# Organization and evolution of sex-biased gene expression in *Drosophila* adult sexual circuits

**DOI:** 10.64898/2026.02.28.708756

**Authors:** Dawn S. Chen, Helena Gifford, Yerbol Z. Kurmangaliyev, Yun Ding

## Abstract

Sexually dimorphic behaviors are widespread across animals. A key open question is how the underlying sexual circuits, which encode these behaviors, evolve to meet sex-specific fitness needs despite being built from largely shared developmental blueprints. Here, we use single-cell transcriptomics, leveraging the sex determination gene *fruitless* (*fru*) as a molecular handle, to examine the transcriptomic landscape and evolution of adult *Drosophila* sexual circuits at cellular resolution. We find that sex-biased gene expression is limited, highly cell-type-specific, and largely species-specific. Species divergence in gene expression is widespread and predominantly coupled between sexes, with sex-specific changes constrained by pleiotropic gene expression across cell types. Furthermore, a strong overlap of differentially expressed genes across sexes, species, and *fru* status points to a common set of genes, particularly neural signaling genes, whose expressions are highly labile across multiple contexts. Finally, transcriptomic divergence between sexes decreases from mid-pupae to adulthood, underscoring the developmentally dynamic nature of sex-biased gene expression. Our results provide fundamental insights into the organization and evolutionary mode of sex-biased gene expression, suggesting that sex-specific adaptation via selective gene programs at localized circuit nodes is a key mechanism that preserves circuit evolvability despite pervasive transcriptomic coupling between sexes.

## Introduction

Sexual behaviors are diverse and essential for reproduction, with each sex often forming a distinct behavioral repertoire to optimize its reproductive fitness. Sexual conflict occurs when sexual behaviors in one sex reduce the fitness in the other sex (Arnqvist and Rowe 2013). One fundamental form of sexual conflict, known as intralocus sexual conflict, arises from the fact that sexes typically share the majority of their genome, such that expression of an allele with positive genetic correlation between sexes may elicit opposite fitness consequences in either sex, thereby driving sexually antagonistic selection (Rice and Chippindale 2001; Bedhomme and Chippindale 2007; Bonduriansky and Chenoweth 2009; van Doorn 2009; Tosto et al. 2023). One way to resolve intralocus sexual conflict is by disrupting the intersexual genetic correlation through the evolution of sex-biased gene expression (Lande 1980; Rice 1984). As such, sex-biased gene expression may represent a transcriptomic resolution and footprint of sexual conflict (Innocenti and Morrow 2010).

In the nervous system, sexual circuits formed from neurons that encode sexual behaviors are the expected hotspot of intralocus sexual conflict. As a result of this conflict over the behavioral fitness optima, sexual circuits exhibit sexual dimorphism in cell types and gene expression that underlie sex differences in behavior. Nonetheless, despite sexually dimorphic nodes, female and male sexual circuits largely derive from common developmental origins, and brain regions and neuron populations responsible for sexual behaviors are often found in both sexes (Cooke et al. 1998; Mank and Rideout 2021). Therefore, at the cellular level, evolutionary changes in one sex may also occur in the other sex due to shared gene regulation that stems from shared developmental origins. As evolutionary changes continue to arise and inflict intralocus sexual conflict, to what extent sex-specific adaptation may emerge to resolve ongoing sexual conflict and which gene programs contribute to such changes remain unknown.

Sex-biased gene expression, as a major mechanism for resolving sexual conflict to permit sex-specific adaptation, has been extensively investigated across a wide range of species. Underscoring the prevalence of sexual conflict and sexually antagonistic selection, whole-animal and whole-tissue transcriptomic studies in mice, *Xenopus*, and *Drosophila* have indicated that the majority of the genomes show sex-biased gene expression (Malone et al. 2006; Yang et al. 2006; Wayne et al. 2007; Ayroles et al. 2009; Innocenti and Morrow 2010). However, whole-animal studies can produce inflated estimates due to the inclusion of gonads, which are highly sexually differentiated tissues whose expression patterns may not be generalizable to other tissues (Stewart et al. 2010). While whole-brain comparisons provide a more direct estimate of sex-biased gene expression in the nervous system, cell-type-specific signals are diluted in a largely isomorphic tissue. Single-cell transcriptomics offer a solution to these confounding factors by affording cell-type-level resolution. Moreover, isolating sexual circuits for single-cell comparisons can prioritize meaningful cellular coverage for the neuronal “battleground” of sexual conflict to better capture cell-type-specific patterns, although this is only feasible in very few systems.

As in many other animals, *Drosophila* species display a suite of rapidly evolving sexually dimorphic behaviors that ranges from courtship, aggression, to locomotion (Markow and O’Grady 2005; Asahina 2018; Jiang et al. 2024). For example, *D. melanogaster* males exhibit an intricate dynamic courtship ritual involving orientation, tapping, waggling, wing vibration (“song”), licking, abdomen quivering, and attempted copulation (Billeter et al. 2006; Pavlou and Goodwin 2013). Females, conversely, respond to these signals by running away or slowing down, as well as female-specific behaviors such as vaginal plate opening or ovipositor extrusion to indicate sexual interests (Aranha and Vasconcelos 2018; Wang et al. 2021). These behaviors are predominantly encoded by neurons expressing the sex determination genes *fruitless (fru)* and/or *doublesex* (*dsx*), two master regulators of sexual differentiation in the nervous system (Billeter et al. 2006; Siwicki and Kravitz 2009; Yamamoto and Koganezawa 2013). Of the two genes, *fru* is more broadly expressed (present in ∼2% of central nervous system (CNS) neurons) and *fru* neurons include most *dsx* neurons in males (Lee et al. 2000; Stockinger et al. 2005; Siwicki and Kravitz 2009; Walsh et al. 2025). The transcripts of *fru* under the P1 promoter are spliced into sex-specific isoforms (Ito et al. 1996; Ryner et al. 1996; Anand et al. 2001). The male isoform *fruM* is translated to full length FruM proteins that function as transcription factors, and the female isoform *fruF* contains an early stop codon and is translated into a truncated protein (Ito et al. 1996; Ryner et al. 1996; Anand et al. 2001; Pan et al. 2025). Abolishing *fru* expression or silencing *fru* neurons largely disrupts courtship in males, while expressing the male-specifc FruM protein in females induces male-typical courtship (Kitamoto 2002; Demir and Dickson 2005; Manoli et al. 2005; Clyne and Miesenböck 2008; von Philipsborn et al. 2011; Chen et al. 2021). Despite known sex differences in some cell types and neuronal morphology, the gross neuroanatomy of *fru* neurons are overall similar between sexes (Kimura et al. 2005; Stockinger et al. 2005; Kimura et al. 2008). The central function of FruM as a master regulator of sexual circuits and the conservation of *fru* splicing (Gailey et al. 2006; Bertossa et al. 2009; Baker et al. 2024) render *fru* a genetic inroad to understanding how sexual circuits evolve under the constraint of largely sex-shared circuit elements and sex-specific fitness outcomes to mediate behavioral evolution.

Here, we uncover the transcriptomic landscape and evolutionary mode of sexual circuits under intralocus sexual conflict at cell-type resolution, employing single-cell RNA sequencing (scRNA-seq) of genetically labelled sexual circuits offered by the molecular handle *fru*, in *D. melanogaster* and *D. yakuba*. This species pair is separated by 7-13 million years (Tamura et al. 2004; Suvorov et al. 2022), and is an emerging comparative model for behavioral and neural evolution (Ding et al. 2019; Coleman et al. 2024; Ye et al. 2024; Walsh et al. 2025). We profiled a total of 58,028 adult cells in females and males in both species, as well as FruM-null mutant males in *D. melanogaster*. Combining these data with published scRNA-seq data of *fru* neurons from an earlier developmental timepoint (Palmateer et al. 2023), we examine the extent, distribution, evolution, and temporal dynamics of dimorphic gene expression in sexual circuits. Our results reject the expectation of pervasive dimorphic gene expression in sexual circuits, and show that sex-biased gene expression is limited, localized, developmentally dynamic, and undergoes rapid evolutionary turnover. Crucially, transcriptomic evolution of sexual circuits is largely coupled between sexes, with sex-specific adaptation constrained by pleiotropic gene expression and preferentially contributed by selective gene programs with flexible expression, particularly neural signaling genes, at localized circuit nodes.

## Results

### Characterization of *fru* neuron clusters by scRNA-seq across sexes, species, and *fru* status

To ensure direct comparison of *fru* circuit elements between species, we generated *fru P1-GAL4* (henceforth *fru-GAL4*) in both *D. melanogaster* and *D. yakuba* with an identical design (Figure S1A), by introducing a GAL4 sequence to the identical location in the first coding exon after the P1 promoter (i.e., the sex-specifically spliced S exon; Figure 1A) using CRISPR/Cas9 genome editing. We tagged the GAL4 with muscle-specific Mhc-DsRed to avoid fluorescence in the nervous system. By driving *UAS-nls::tdTomato* with *fru-GAL4* and isolating tdTomato+ cells using fluorescence-activated cell sorting (FACS), we generated single-cell transcriptomes of male and female *fru* neurons in both species. In addition, to understand the contribution of genes regulated by FruM in species and sex differences, we generated single-cell transcriptomes of the *fru* neurons of FruM-null males in *D. melanogaster* (Figure S1B).

**Figure 1.**
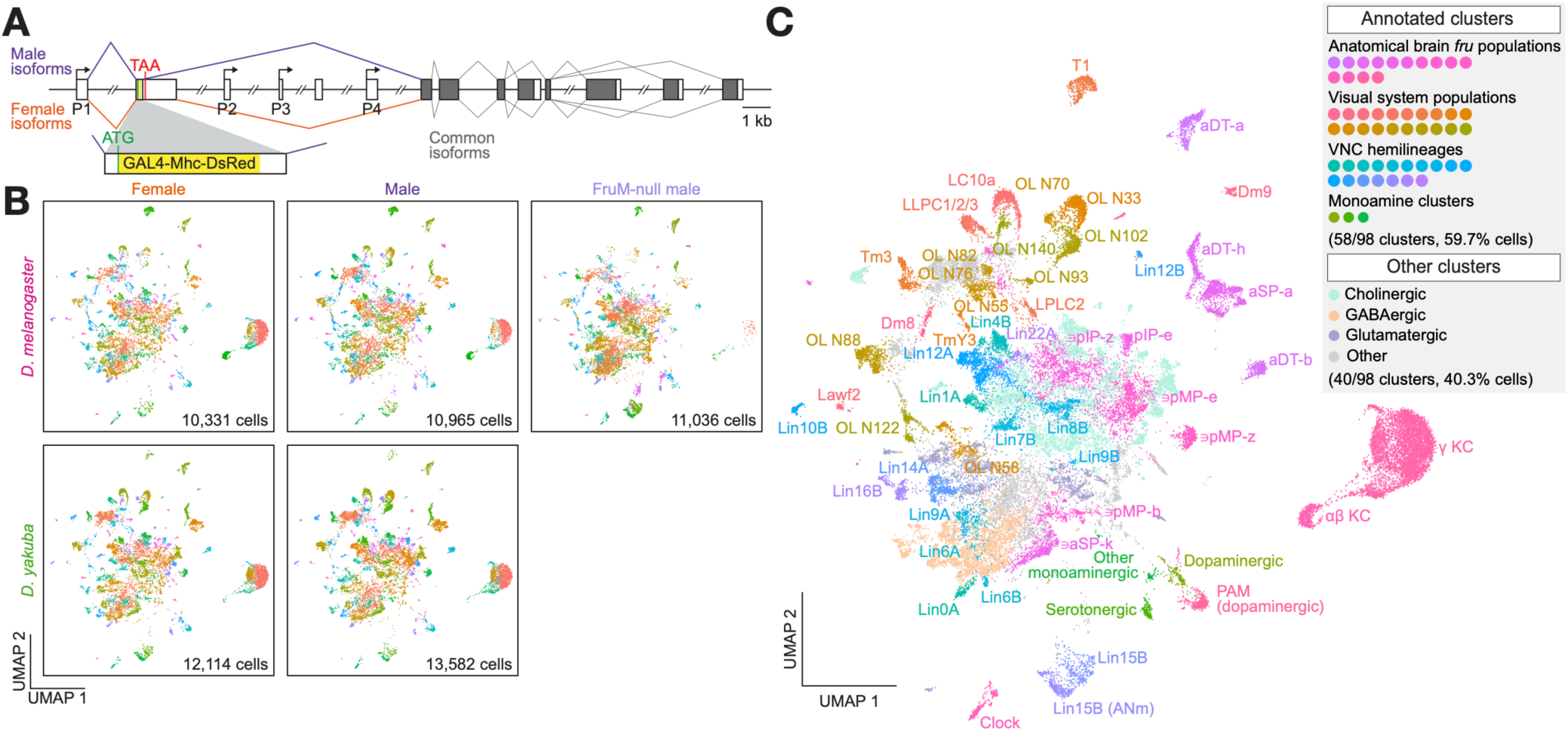
Sample integration and *fru* neurons annotation. **(A)** Insertion of the *T2A-GAL4-Mhc-DsRed* transgene at the endogenous *fru* locus in *D. melanogaster and D. yakuba.* Male-specific, female-specific, and sex-common splicing are shown in purple, orange, and gray, respectively. Inset shows exon 2 with male-specific splicing. **(B)** Uniform Manifold Approximation and Projection (UMAP) of each of the five sample types. Number of cells included in each sample type are shown in the bottom right corner of each panel. **(C)** UMAP ofclusters annotated by anatomical brain population, visual system population, VNC hemilineage, or fast-acting neurotransmitter identity. OL: optic lobe; ANm: abdominal neuromere.

After filtering for 1:1 orthologs between *D. melanogaster* and *D. yakuba* and additional quality filters (see Methods), we obtained 58,028 high-quality cells from the 5 sample types with 2 replicates each (mean±SD: 5,802±1,006 cells/sample). Replicates of each sample type were highly congruent (Figure S1C; quality metrics in Figure S1D, Supplementary Table 1). Our data represents a total of 9.28× cellular coverage among the four wildtype sample types (see Methods). We note that the commonly cited estimate of ∼2,000 *fru* neurons in the CNS likely represents an underestimation, as the imaging-based cell counts excluded the mushroom body and the visual system (Stockinger et al. 2005). Instead, our data estimates the presence of over 4,000 *fru* neurons in the CNS (see Methods), comparable to the connectome-based estimate (Berg et al. 2025), constituting 2-3% of the entire CNS.

Our data consisted of 102 transcriptomically defined clusters, which reflected a mostly 1:1 relationship to major brain anatomical populations and VNC hemilineages (Figure S2A). Clustering resolution had little impact on overall group assignment, and increasing resolution mostly subdivided larger clusters (Figure S1E). All but the smallest cluster were represented in all sample types (cluster 101 was only found in *D. melanogaster*). 99.4% of all cells expressed at least one of the neuronal markers *elav, brp, Syt1*, or *CadN* (McLaughlin et al. 2021). The detection rate of *fru* in all cells (81.4%) was on par with the detection rate of individual neuronal maker genes (76.5%-97.8%) (Supplementary Table 2), validating the reliability of isolating *fru* neurons using genetic labeling. Out of caution, we removed 4 clusters in which <90% of cells expressed any of the neuronal marker genes *elav, brp, Syt1,* or *CadN* or <50% of cells were *fru+* from further analysis (Figure S1F-G). Less than 0.1% of cells expressed the glia marker *repo* (Freeman 2015). As a further assessment of data quality, we generated bulk RNA-seq data on FACS-sorted *D. melanogaster* and *D. yakuba* male *fru* neurons, and compared the transcript counts to those aggregated over each sample type (pseudobulked) in our single-cell data. Gene expression levels correlated well between methods for each sample type (Figure S1H), lending further confidence in the quality of our single-cell data.

As expected, *fru* neurons were highly transcriptomically heterogeneous, but homologous clusters integrated well between species and sexes in the UMAP space (Figure 1B, Figure S2B-C). Using known molecular markers, we annotated each cell by anatomical region (brain and ventral nerve cord (VNC); Figure S2D), VNC segment Hox gene expression (Figure S2E) fast-acting neurotransmitter expression (Figure S2F), and monoamine expression (Figure S2G). Leveraging published transcriptomic atlases (Kurmangaliyev et al. 2020; Naidu et al. 2020; Özel et al. 2021; Palmateer et al. 2023; Cachero et al. 2025; Allen et al. 2026) and incorporating these cell-level annotations, we further annotated 34 brain anatomical populations, 17 VNC hemilineages, and 3 monoaminergic clusters, accounting for 58/98 of the clusters and 59.7% of the cells in the dataset (Figure 1C, Supplementary Table 3), providing a most comprehensive annotation of *fru* neurons based on scRNA-seq data to date.

### *fru* clusters defined by scRNA-seq capture known major sex differences in cell types and are otherwise largely shared between sexes

Given the central role of *fru* neurons in male sexual behaviors, one might expect sexual dimorphism across various dimensions that influence neuronal function, such as cell type abundance, overall transcriptomic profiles, and differential expression of specific genes. Indeed, our data recapitulated previously reported male-biased or male-specific *dsx*+ cell types pC1, pC2l, pC2m, and TN1 (the *fru*+ cells of which were captured in our data), and the *fru+* cell type mAL (Figure 2A-B, Figure S3). Each of these cell types occupied either defined subclusters within a *fru* cluster or the entire cluster. For example, pC1 neurons are known for their sexually dimorphic abundance (48-70 neurons/hemisphere in males (Kimura et al. 2015; Nojima et al. 2021; Walsh et al. 2025) and 6-9 neurons/hemisphere in females (Nojima et al. 2021; Walsh et al. 2025)) and roles in sensory integration and regulation of sexual behaviors in both sexes (Cachero et al. 2010; von Philipsborn et al. 2011; Zhou et al. 2014; Hoopfer et al. 2015; Koganezawa et al. 2016; Deutsch et al. 2020; Wang et al. 2020). In our data, *fru*+ pC1 cells occupied two subclusters of cluster 7, and sexual dimorphism manifested in males having more cells in these subclusters and sex-specific expression of *dsx* in both species (Figure 2A). We note that females had *fru+* counterparts to pC1, pC2l, and TN1 that occupied the same regions in the respective UMAP spaces, but they lacked *dsx* expression (Figure 2A-B, Figure S3B).

**Figure 2.**
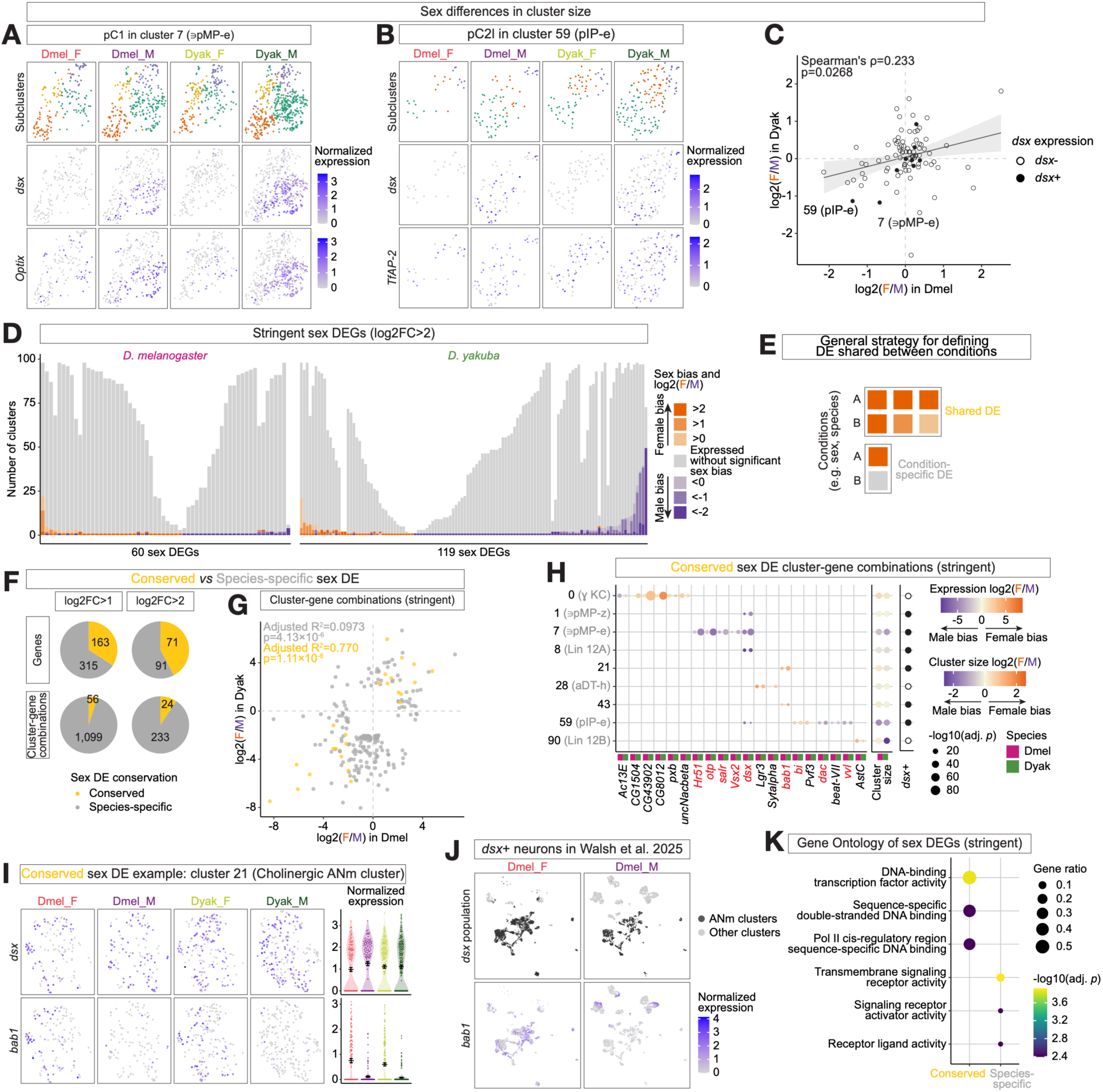
Sex differences are limited and largely species-specific. **(A)** Cluster 7 plotted in its own Uniform Manifold Approximation and Projection (UMAP) space, separated by sample type. The top row shows cells color-coded by their subcluster identity, and the middle and bottom rows show cells color-coded by normalized expression of pC1 marker genes *dsx* and *Optix,* respectively. Color scales for their respective genes are shown to the right. **(B)** Cluster 59 plotted in its own UMAP space, separated by sample type. The top row shows cells color-coded by their subcluster identity, and the middle and bottom rows show cells color-coded by normalized expression of pC2l marker genes *dsx* and *TfAP-2,* respectively. Color scales for their respective genes are shown to the right. **(C)** Correlation of Iog2-transformed ratio of female cells to male cells in each cluster in *D. melanogaster versus D. yakuba.* Dots are color-coded by each cluster’s dsx expression status. Clusters 7 and 59 are labeled. Regression line and 95% confidence interval are shown behind data points in gray. **(D)** Number ofclusters in which each *D. melanogaster* (left) and *D. yakuba* (right) sex differentially expressed gene (DEG) is expressed and sex DE in its respective species. Clusters in which each sex DEG is expressed without significant sex bias are shown in gray, and clusters with sex-biased expression are shown in shades of orange (female-biased) or purple (male-biased) based on their Iog2-transformed ratio of aggregated female to male gene expression (log2FC). **(E)** Diagram illustrating the general strategy for defining DE shared between conditions such as sexes or species. Darker shades of orange denote higher log2FC, and gray denotes expression without significant species bias. **(F)** Pie charts showing the number of conserved (yellow) and species-specific (gray) sex DE genes (top row) and cluster-gene combinations (bottom row) under two log2FC thresholds. Number of elements in each sector are shown in the respective sector. **(G)** Log2FC of sex DE cluster-gene combinations in *D. melanogaster versus D. yakuba.* Dots are color-coded by whether each cluster-gene combination is conserved (yellow) or species-specific (gray). **(H)** Log2FC and -log 10-transformed adjusted p-value (-Iog10(adj. p)) of conserved sex DE cluster-gene combinations in both species. Log2-transformed ratio of female cells to male cells and *dsx* expression status for each cluster are shown to the right. Size scale of-Iog10(adj. p) and color scales of log2FC are shown next to the plot. Magenta and green rectangles in the margin of each column denote *D. melanogaster* and *D. yakuba,* respectively. Transcription factors are highlighted in red. **(I)** Cluster 21 plotted in its own UMAP space, separated by sample type. The top and bottom rows show cells color-coded by normalized *dsx* and *bab1* expression, respectively. Normalized expression of each gene in each cell, grouped by sample type, are shown to the right. Error bars show mean±SEM. **(J)** *dsx+* neurons in Walsh et al. 2025 colored by whether they are abdominal neuromere (ANm) clusters (top) and *bab1* normalized expression (bottom), separated by sex. **(K)** Molecular function Gene Ontology term enrichment of genes in conserved *versus* species-specific sex DE cluster-gene combinations, showing enrichment terms with adjusted p-value<0.01 and gene ratio>0.03. Color scale of-Iog10(adj. p) and size scale of gene ratio are shown next to the plot. KC: Kenyon cells; ANm: abdominal neuromere; Dmel: *D. melanogaster,* Dyak: *D. yakuba;* F: female; M: male.

Beyond recapitulating these known sex differences, we found that sex differences in cluster size were limited overall. We first normalized cell counts in *D. melanogaster* based on empirical estimates in the respective sex (Stockinger et al. 2005), and in *D. yakuba* under the assumption that the median cluster was conserved in size between the species pair. After normalization, 23 of the 98 clusters met an abundance difference threshold of log2 fold-change (FC)>1 in either species and only 3 met a stringent threshold of log2FC>2 in either species (Figure 2C). The clusters with the strongest conserved male biases were the *dsx*+ pC1-containing cluster 7 (annotated as containing (∋) pMP-e) and pC2l-containing cluster 59 (pIP-e). Nonetheless, no cluster had statistically significantly sex-biased abundance at FDR<0.2 using the Bayesian scCODA model for compositional data (Büttner et al. 2021). Overall, our data captures known sexual dimorphism in cell types, and at the same time did not reveal extensive major sex differences in cluster size, demonstrating that sexual dimorphism in cell types is highly localized even within the *fru*-expression neurons.

### Sex differences in gene expression are overall limited and highly cluster-specific

Contrary to an expectation of broad sex-biased gene expression underlying sex differences in *fru* circuit function, genes differentially expressed (DE) between sexes (i.e., sex DEGs) in adult *fru* neurons appeared limited: 191 genes in *D. melanogaster* and 371 genes in *D. yakuba* (2.97% and 5.77% of genes expressed in *fru* neurons) were DE between sexes (log2FC>1 and adjusted p-value<0.05 in any cluster), and only 60 and 119 genes (0.933% and 1.85%), respectively, further met the stringent sex DEG threshold of log2FC>2. There was a similar amount of female- and male-biased sex DE cluster-gene combinations in *D. melanogaster* and more male-biased combinations in *D. yakuba* (Figure S4A). On average, each cluster had 8.37±11.6 sex DEGs at logFC>1 or 3.09±3.65 at log2FC>2. Notably, the clusters with the strongest conserved male-biased abundance (clusters 7 and 59) were among the clusters with the most sex DEGs (30.5±20.4 at logFC>1; 10.5±5.74 at log2FC>2).

Sex DEGs were highly cluster-specific. Defining a gene with >10% detection rate in a cluster as being expressed, we found that each sex DEG was only DE in one to a few clusters (2.49±4.43 clusters at log2FC>1; 1.60±3.81 clusters at log2FC>2) despite their generally broad expression (74.8±30.0 clusters at log2FC>1; 56.0±33.7 clusters at log2FC>2) among *fru* neurons (Figure 2D, Figure S4B). A large proportion of these sex DEGs were only differentially expressed in a single cluster regardless of fold-change thresholds (77.0%-88.3% in *D. melanogaster*; 66.8%-84.9% in *D. yakuba*). Of note, genes that were sex DE in multiple clusters mostly exhibited concordant directions of sex bias across clusters (Figure 2D, Figure S4B-C), suggestive of shared sex-specific gene regulation among cell types. Together, these results indicate two complementary themes of sexually dimorphic gene expression: a strong cell-type-specificity in the regulation of sex-biased gene expression, and regulatory pleiotropy (i.e., shared gene regulatory mechanisms across cell types) in imposing a largely consistent sex bias across DE clusters. In summary, sex DEGs are limited and highly localized, and adult *fru* neurons lack extensive transcriptional remodeling between sexes.

### Evolutionary turnover and conservation of sex-biased gene expression

When comparing sex DEGs between species, we found very little overlap of sex DEGs between *D. melanogaster* and *D. yakuba*. To mitigate underestimation arising from threshold sensitivity, we focused on sex DEGs (log2FC>2 or >1) in either species and defined conserved ones as those with p<0.05 in the other species without requiring a further fold-change threshold (Figure 2E). Even with this approach, the majority of sex DEGs were species-specific, and sex DE cluster-gene combinations were overwhelmingly species-specific (Figure 2F, Figure S4D, Figure S5), suggesting that while a set of genes might be common substrates of sex-biased gene expression, the cellular context of such sex-bias was highly evolvable. Moreover, the log2FC of species-specific sex DE cluster-gene combinations were poorly correlated between species (Figure 2G), further underscoring the rapid evolutionary turnover of sex-biased gene expression.

We focused on the 24 conserved sex DE cluster-gene combinations, corresponding to 20 genes, under the stringent log2FC>2 threshold to understand their function (Figure 2H; conserved sex DEGs with 1<log2FC<2 are shown in Figure S4E). Overall, these conserved DEGs were primarily found in *dsx*+ clusters, with many male-biased DEGs reflecting sex differences in the relative abundance of cell types within the cluster. For example, *dsx* was differentially expressed in a male-biased manner in clusters 1 (∋pIP-z), 7 (∋pMP-e), 8 (Lin12A), and 59 (pIP-e), which contained neurons of pC2m, pC1, TN1, and pC2l, respectively, the four aforementioned male-specific or -biased *dsx*+ cell populations (Kimura et al. 2015; Nojima et al. 2021; Walsh et al. 2025). Additionally, all other genes with male-biased expression in cluster 7 (*Hr51*, *otp*, *salr*, and *Vsx2*) were transcription factors and strong marker genes for the male-biased pC1 cell types (Walsh et al. 2025; Allen et al. 2026).

This dataset also recapitulated the well-characterized female-biased expression of *Lgr3* in aDT-h neurons (cluster 28) (Meissner et al. 2016; Allen et al. 2026) and discovered previously unknown sex DEGs in the CNS. Most notably, we identified female-biased gene expression of the transcription factor *bab1* in the cholinergic cluster 21 and the GABAergic cluster 43 (Figure 2I, Figure S4F), both of which are *dsx*+ abdominal neuromere clusters (*abd-A+ Abd-B+*) in the VNC. Since these two clusters were not sex-biased in cluster size or *dsx* expression, and *bab1* was broadly expressed throughout the cluster UMAP spaces in females, the DE likely reflected a sex difference in gene regulation rather than cell type abundance. Further inspection of a recently available scRNA-seq dataset of *dsx*+ neurons in *D. melanogaster* (Walsh et al. 2025) showed that *bab1* was widely expressed in a female-specific manner in *dsx+* abdominal neuromere neurons (cells that expressed *abd-A* and/or *Abd-B*), but not in other *dsx+* populations (Figure 2J). A similar female-specific expression of *bab1* in *dsx+ Abd-B+* cells has been reported in abdominal epidermis cells, where Abd-B cooperates with the female-specific Dsx isoform to activate *bab1* expression in females and produce sexually dimorphic abdominal pigmentation (Kopp et al. 2000; Williams et al. 2008). These parallel observations may suggest a common genetic pathway between morphological and behavioral traits in integrating segment and sex identities to deploy dimorphic regulatory programs. Alternatively, because *bab1* is also a temporal factor whose expression correlates with birth order (Allen et al. 2025; Cachero et al. 2025; Elkahlah et al. 2025), its female-biased expression may result from temporal differences in the birth of female and male *dsx+* abdominal neurons during neurogenesis.

Gene Ontology (GO) enrichment analysis revealed a major difference between conserved and species-specific sex DEGs: the former were strongly enriched for transcription factors, and the latter for signaling receptor activity (Figure 2K). Genes in the latter group included neuropeptides such as *NPF*, *Dh44*, and *AstA*, and G protein-coupled receptors (GPCRs) such as *Dop2R*, *SPR*, *CCHa1-R*, and *Dh31-R*, many of which are known to regulate sex differences (Yapici et al. 2008; Kim et al. 2013; Jiang et al. 2023; Kim et al. 2024; Biswas and Rideout 2025). These results might suggest that sexual dimorphism is established by a core set of conserved transcription factors and potentially evolutionarily labile downstream gene batteries to encode sex differences in neuronal function.

### Species divergence in cluster size is correlated between the sexes

We found a large number of clusters with potential species divergence in cluster size: 20 out of 98 (20.4%) clusters diverged in abundance in either or both sexes at the scCODA FDR<0.2 level (six clusters with FDR<0.05), and 10 of them with log2 fold-change>2 (Figure 3A). Species differences in cluster size were strongly coupled between sexes, with females and males showing highly correlated fold-changes in cluster size between species (Figure 3A). The same patterns were also observed at the subcluster level (Figure S6A). We note that species differences in cluster size observed in *fru* neurons could arise from the cellular expansion or contraction of a transcriptomically defined *fru* cluster, or the gain or loss of *fru* expression. Given that even fine-grained neuronal cell types are largely conserved between *D. melanogaster* and *D. yakuba* (Walsh et al. 2025), the cluster size differences may simply result from changes of *fru* expression that affect both sexes, reflecting a lack of sex-specific regulation of *fru* transcription in evolution. This is conceivable from the perspective of intralocus sexual conflict: the sex-specific splicing of *fru* to produce a male-specific transcription factor means that *fru* gene expression evolution is presumably functionally consequential only in males, such that potential sexual conflict that would otherwise arise from sex-shared transcriptional regulation is effectively bypassed. While species differences in *fru* clusters’ cell abundance may be common, none of the nine *dsx*+ clusters showed significant species differences in either sex, and *dsx*+ subclusters were also overall conserved in size (Figure 3A, Figure S6A). This is consistent with our earlier finding that *fru* expression is highly conserved among *dsx*+ neurons (Walsh et al. 2025), and raises the possibility of *fru* expression being evolutionarily more labile outside of *dsx*+ neurons.

**Figure 3.**
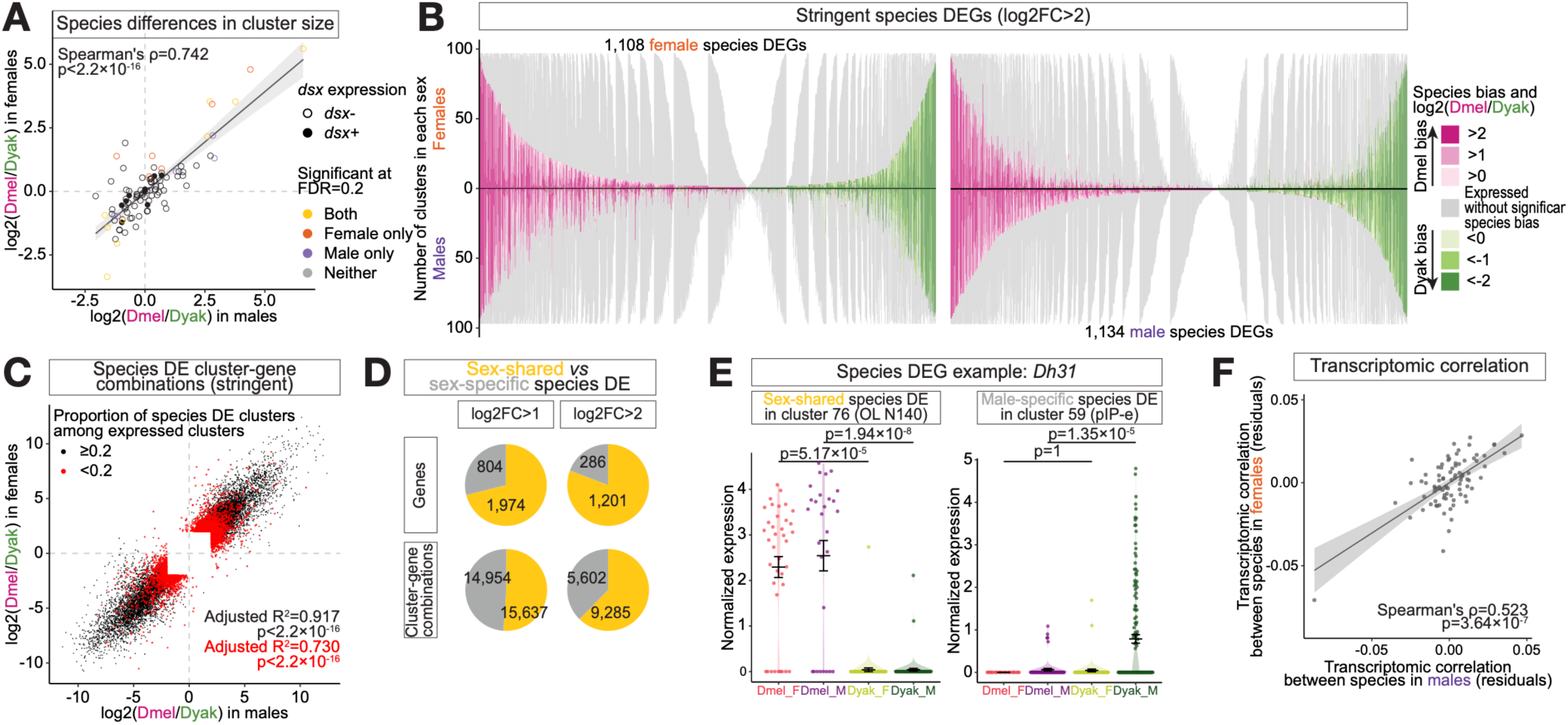
Species differences are prevalent and extensively sex-coupled. **(A)** Correlation of Iog2-transformed ratio of D. *melanogaster* cells to *D. yakuba* cells in each cluster in females *versus* males. Filled dots denote *dsx+* clusters, and empty dots denote dsx-clusters. Dots are color-coded by the sex(es) in which their species bias is statistically significant at False discovery rate (FDR)=0.2. Regression line and 95% confidence interval are shown behind data points in gray. **(B)** Comparison of female (left) and male (right) species differentially expressed gene (DEG) expression in females (above x-axis) versus males (below x-axis). Clusters in which the species DEG is expressed without significant species bias are shown in gray, and clusters with species-biased expression are shown in shades of magenta (D. *melanogaster-blased)* or green (D. yakuba-biased) based on their Iog2-transformed ratio of aggregated *D. melanogaster to D. yakuba* gene expression (log2FC). **(C)** Log2FC of species differentially expressed (DE) cluster-gene combinations in males *versus* females. Dots are color-coded by whether the gene in each cluster-gene combination is species DE in >20% (black; more broadly species DE) or <20% (red; highly cluster-specific species DE) of the clusters in which it is expressed. **(D)** Pie charts showing the number of sex-shared (yellow) and sex-specific (gray) species DE genes (top row) and cluster-gene combinations (bottom row) under two log2FC thresholds. Number of elements in each sector are shown in the respective sector. **(E)** Normalized expression *Dh31* in a cluster with sex-shared species DE and one with male-specific species DE. Cells are color-coded and grouped by sample type. Error bars show mean±SEM. Statistical significance is tested with Seurat’s “FindMarkers” function with the MAST method. OL: optic lobe. **(F)** Correlation of inter-species Kendall’s tau transcriptomic correlation in males *versus* females. In each sex, inter-species Kendall’s tau transcriptomic correlation are calculated for each cluster and the confounding variables (normalized cluster size, inter-replicate Kendall’s tau transcriptomic correlation, and number of genes used to calculate transcriptomic correlation) are regressed out to obtain the residuals. Regression line and 95% confidence interval are shown behind data points in gray. Dmel: *D. melanogaster,* Dyak: *D. yakuba’,* F: female; M: male.

### Transcriptomic evolution is strongly coupled between sexes

A major hallmark of sexual conflict is the evolution of sex-biased gene expression, which decouples evolutionary changes in gene expression between sexes to allow sex-specific adaptation (Mank 2017; Tosto et al. 2023). Here, we identified DEGs between *D. melanogaster* and *D. yakuba* for each sex and examined the extent to which they were shared between sexes. Gene expression differences between the two species appeared widespread: applying the same thresholds as defining sex DEGs (log2FC>2 and adjusted p<0.05 in any cluster), we found 1,108 and 1,134 species DEGs (17.2% and 17.6% of genes expressed in *fru* neurons) in females and males, respectively. In each sex, most identified species DEGs were DE in only a small subset of clusters in which they were expressed (77.3% and 72.8% of female and male species DEGs, respectively, were species DE in <20% of the clusters they were expressed in), whereas a smaller fraction were DE across many clusters (9.48% and 12.0% of female and male species DEGs, respectively, were species DE in >50% of the clusters they were expressed in). The expression breadth and species bias in gene expression were closely mirrored between sexes (Figure 3B, Figure S7). Consistently, species DEGs’ gene expression differences between species (i.e., log2FC) were highly correlated between sexes (Figure 3C).

Furthermore, species DE genes and cluster-gene combinations were both well-shared between sexes and concordant in the direction of their species bias, and this finding was robust at different stringencies of DEG analyses (Figure 3D; example shown in Figure 3E). Of all cluster-gene combinations tested in both sexes, none showed discordance in species bias (i.e., none were *D. melanogaster*-biased in one sex and *D. yakuba*-biased in the other sex). These results demonstrated that the widespread species DEGs were predominantly shared between sexes at the levels of gene, gene-cluster combination, and the direction of species-bias. In addition to DEGs, we also assessed species divergence in overall transcriptomic patterns for each cluster in both sexes using Kendall’s tau correlation coefficient, a non-parametric statistic that measures the ordinal association of gene expression between pairs of samples. As a proof of principle, and consistent with the relative abundance of sex DEGs versus species DEGs, transcriptomic correlation was much lower in species comparisons than in sex comparisons (Figure S6B). After accounting for confounding variables that influenced transcriptomic correlations (see Methods), we found a moderate association of inter-species transcriptomic correlation between sexes (Figure 3F).

We acknowledge that cross-species transcriptomic comparisons have important caveats, including potential artifacts caused by sequence divergence and gene annotation errors. These issues may be more pronounced in single-cell RNA-seq than in bulk RNA-seq, because many single-cell platforms preferentially capture the 3’ ends of transcripts, which are particularly sensitive to annotation errors. Such artifacts could generate false-positive DEGs detected in both sexes, thereby inflating the apparent degree of evolutionary coupling between sexes. We attempted to address this issue in two ways. First, using the bulk RNA-seq data of *fru* neurons in males of both species, we validated that the species DEGs identified by scRNA-seq were highly congruent with results of bulk sequencing: the number of clusters that were DE for each species DEG in the scRNA-seq data was predictive of the log2FC between species in the bulk RNA-seq data (Figure S6C). The direction of species bias was also consistent between methods (Figure S6D). Second, we argue that such artifacts would be expected to manifest broadly across clusters. When restricting the analysis to species DEGs that were DE in fewer than 20% of clusters in which the gene was expressed, species differences in gene expression remained strongly correlated between sexes (Figure 3C). Therefore, while we expect some false positives, they are insufficient to drive the pattern of pervasive coupling of cell-type-specific gene expression changes between sexes.

Together, these observations suggest that the evolution of gene expression, even within the most sexually dimorphic part of the nervous system, typically involves high gene expression correlation between sexes, and that sexual conflict does not drive a widespread sex-specific mode of gene expression evolution.

### Factors shaping gene expression evolution

#### Pleiotropic gene expression constrains sex-specific adaptation

Because the sex DEG analysis indicated shared regulatory mechanisms underlying sex-biased expression across cell types, we asked whether the strong transcriptomic coupling between sexes may partially reflect regulatory constraints that limit the evolution of sex-specific expression. Pleiotropic gene expression can constrain the breakdown of intersexual genetic correlation and therefore impose limits on the extent to which sexual dimorphism can evolve in a cell-type-specific manner (Ellegren and Parsch 2007). We calculated the sex-specificity of each species DEG, defined as the proportion of its species DE clusters that were sex-specific (Figure 4A). We found a strong negative correlation between sex-specificity and expression breadth: the more sex-specific a species DEG was, the more narrowly it was expressed (Figure 4B), suggesting that sex-specific gene expression change could be constrained by pleiotropic gene expression.

**Figure 4.**
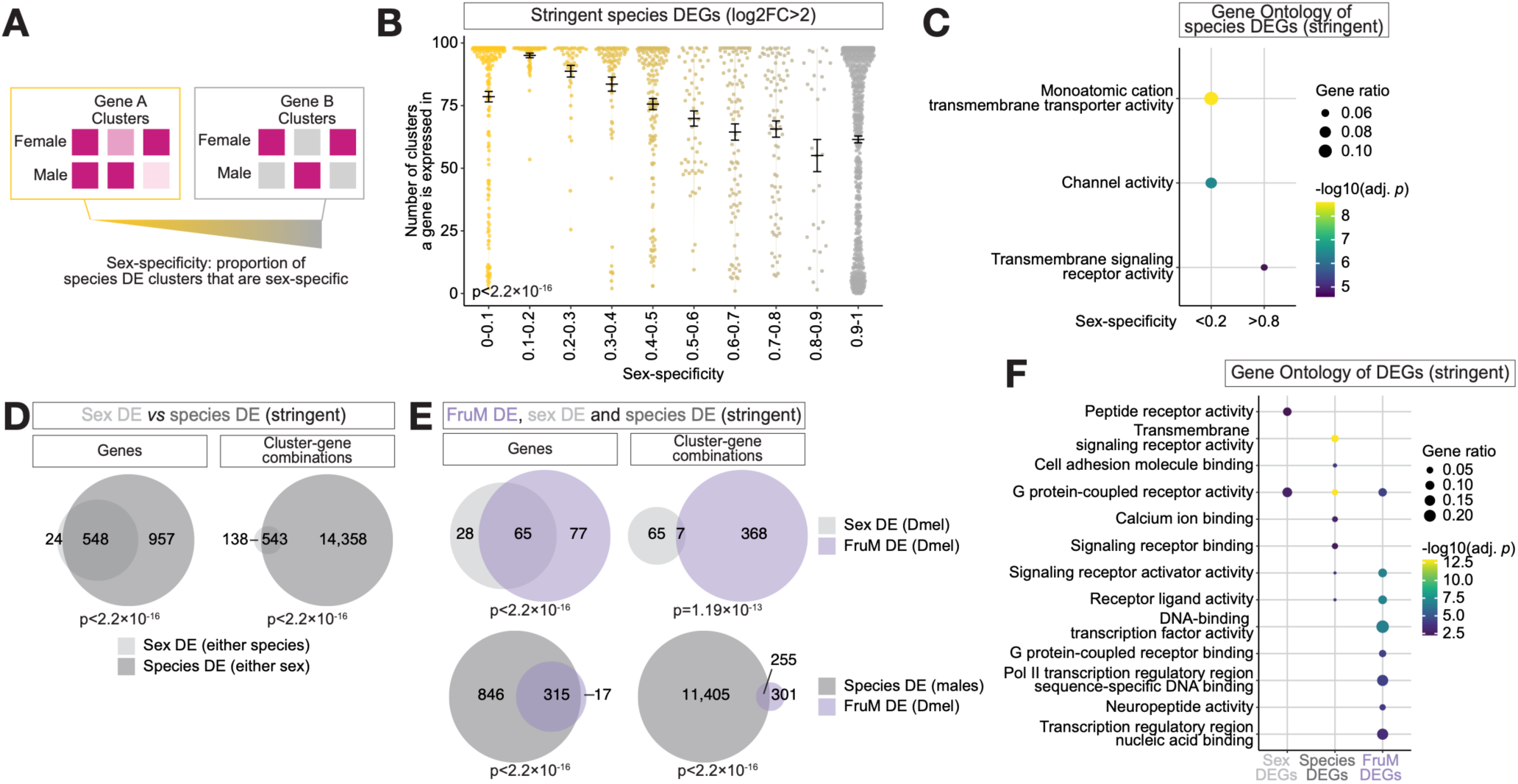
Factors shaping gene expression evolution. **(A)** Diagram illustrating how sex-specificity is calculated for each species differentially expressed gene (DEG). Darker shades of magenta denote higher log2FC, and gray denotes expression without significant species bias. **(B)** Number of clusters that each species DEG is expressed in, with species DEGs grouped by sex-specificity. Error bars show mean±SEM. Statistical significance is tested with ANOVAon a linear model. **(C)** Molecular function Gene Ontology term enrichment of species DEGs with sex-specificity <0.2 or >0.8, showing enrichment terms with adjusted p-value<0.01 and gene ratio>0.03. Color scale of -Iog10(adj. p) and size scale of gene ratio are shown next to the plot. **(D-E)** Area-proportional Euler diagrams showing the number of genes and cluster-gene combinations DE between sexes (sex DE) and between species (species DE) **(D)**, and those DE between sexes in *D. melanogaster (sex* DE), between species in males (species DE), and between FruM-null males and wildtype males (FruM DE) **(E)**. Statistical significance of overlaps are calculated based on the hypergeometric distribution and shown below each Euler diagram. **(F)** Molecular function Gene Ontology term enrichment of sex DEGs, species DEGs, and FruM DEGs with adjusted p-value<0.01 and gene ratio>0.03. Color scale of-Iog10(adj. p) and size scale of gene ratio are shown next to the plot.

#### Different functional categories underlie sex-specific versus sex-shared expression evolution

To understand if genes that evolved species-biased expression with high *versus* low sex-specificity were functionally different, we performed GO enrichment for genes on either end of the sex-specificity continuum (Figure 4C). We found that genes with predominantly sex-shared species DE were enriched for basic cellular functions such as transmembrane transporter/ channel activity, but highly sex-specific genes were enriched for signaling receptor activity and included mostly GPCRs. The latter was reminiscent of the genes in species-specific sex DE cluster-gene combinations also being enriched for receptor activity (Figure 2K), suggesting that a group of genes involved in neural signaling may be particularly poised to evolve gene expression in a sex-specific manner.

#### Sexual dimorphism versus species differences in gene expression

Consistent with the limited overlap of sex DE between species, sex DEGs were predominantly species DEGs: over 95.7% of sex DEGs and over 72.0% of sex DE cluster-gene combinations were also species DE (Figure 4D, Figure S8A), further underscoring the evolutionary lability of sex DEGs in gene expression. We note that rapid evolutionary turnover is a hallmark of sex DEGs that has been observed across many species and tissues, including *Drosophila* brain inferred by bulk RNA-seq (Zhang et al. 2007; Harrison et al. 2015; Catalán et al. 2018; Khodursky et al. 2020; Rodríguez-Montes et al. 2023; Bontonou et al. 2024; Xie et al. 2024), with many possible underlying causes such as sexual selection, sexually antagonistic selection, and relaxed constraints (Ranz et al. 2003; Ellegren and Parsch 2007; Müller et al. 2012).

#### Genes downstream of fru

How sex differentiation mechanisms such as sex determination genes and sex steroid hormone pathways contribute to sex differences in transcriptomic patterns remains poorly understood. Here, leveraging the *fru-GAL4* allele’s characteristic as a functionally null allele, we generated the first single-cell transcriptome of FruM-null males in *D. melanogaster* to examine the role of FruM in organizing sex-biased gene expression at cellular resolution. This dataset also allowed us to test whether genes responsive to FruM function were more likely to contribute to species-level differences.

We found that the majority of sex DEGs were also FruM DEGs, albeit often in different clusters (Figure 4E, Figure S8B). In addition, the transcriptomic profile of FruM-null males did not simply reflect a feminized state of the male transcriptome: FruM-null male samples formed a distinct cluster from wildtype *D. melanogaster* male and female samples in principle component (PC) space (Figure S8C, Figure S9A), and were transcriptomically more distant to wildtypes males than wildtype females were at both the sample and cluster levels (Figure S8D, S9B). The absence of functional FruM proteins may lead to dysregulation of genes downstream of FruM as well as cell type differences (Kimura et al. 2005), the latter of which is exemplified by the expected reduction of mAL neurons (cluster 52) in the FruM-null samples (Figure S8E). On a global level, however, the dramatic transcriptomic differences were not strongly driven by changes in cell type composition between wildtype and FruM-null males, as the aforementioned results were robust to the exclusion of 17 clusters with significant (scCODA FDR<0.2) and strong (among the top 20 clusters with the largest fold-changes) size differences (Figure S8F-G). Interestingly, this dramatic transcriptomic remodeling in FruM-null males resembles findings in mice, where estrogen receptor-expressing neurons in some brain regions encoding sex-typical social behaviors express more DEGs between different female sex hormonal states than between sexes (Knoedler et al. 2024). These results suggest a profound function of key sexual differentiation molecules in regulating the transcriptomic state of sexual circuits rather than a simple binary sex switch.

We then asked if genes regulated by FruM function were also overrepresented among species DEGs. Over 90% of FruM DEGs were species DEGs in males regardless of the stringency threshold, and over 40% of the DE cluster-gene combinations were shared (Figure 4E, Figure S8H). Comparing the functional categories of sex DEGs, species DEGs, and FruM DEGs, we found that the GO term of GPCR activity was enriched in all three groups (Figure 4F). GPCRs did not exhibit higher expression variability between replicates than the genome average (Figure S8I), which argues against flexible expression of GPCRs as an artifact of noisy expression, and instead lends support for GPCRs being unique in their sex-biased expression, expression evolvability, and sensitivity to FruM.

### Reduction of sex differences from mid-pupae to adulthood

A previous study on *D. melanogaster* mid-pupal stage *fru* neurons revealed substantial dimorphic gene expression, albeit under more lenient thresholds (log2FC>0.25, adjusted p-value<0.05, and detection rate>25% (Palmateer et al. 2023)). We asked if this difference was due to differences in methodology or reflected developmental stage-dependent regulation of sex-biased gene expression. The pupal data collected in the previous study and the *D. melanogaster* adult data presented in our study were too different to be integrated or accurately matched (see Methods), so we kept both datasets separate and reanalyzed the pupal data using a consistent methodology. At the sample level, pupal samples showed clear separation between sexes based on transcriptomic similarity, while adult samples did not (Figure 5A, Figure S9B). A similar pattern was also observed in a bulk RNA-seq study on non-mushroom body *fru* neurons (Brovkina et al. 2021). Consistent with a clear transcriptomic separation by sex in pupal stage but not adulthood, pupal clusters also showed lower overall transcriptomic correlation between sexes than their adult counterparts (Figure 5B).

**Figure 5.**
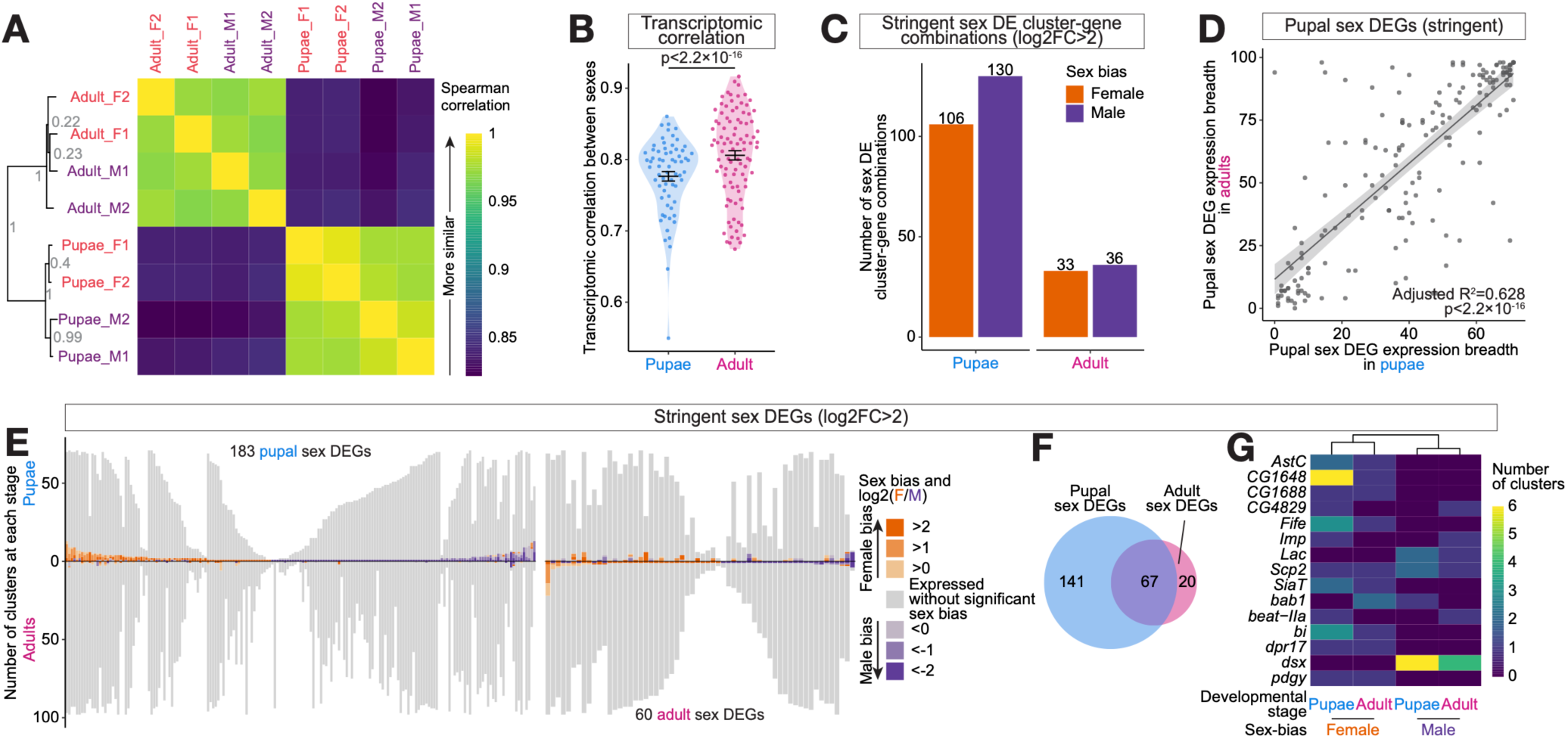
Sex differences of *fru* neuron transcriptomes reduce from pupae to adulthood. **(A)** Left: neighbor-joining tree constructed based on pairwise sample-level Spearman correlation calculated using the log-normalized aggregated gene expression of the 2,000 most variable genes across the eight samples. Numbers next to each node show the proportion of draws (out of 1000 random draws of 1000 genes each, without replacement) that support each branch. Right: Heatmap showing Spearman correlation between each pair of samples. Color scale is shown on the right. Adult_F1, Adult_F2, Adult_M1, and Adult_M2 correspond to melF1, melF1, melM1, and melM2, respectively. F: female; M: male. **(B)** Inter-sex Kendall’s tau transcriptomic correlation of each cluster in *D. melanogaster pupae* and adults. Error bars show mean±SEM. Statistical significance is tested with ANOVA on a linear model that also accounts for cluster size (log-transformed proportion of each sample), inter-replicate Kendall’s tau transcriptomic correlation, and number of genes used to calculate transcriptomic correlation. **(C)** Number of sex differentially expressed (DE) cluster-gene combinations in pupae and adults, color-coded by sex bias. Number of genes in each category are shown above each bar. **(D)** Number of clusters in pupae *versus* adults that each pupal sex DEG is expressed in. Regression line and 95% confidence interval are shown behind data points in gray. **(E)** Comparison of pupal (left) and adult (right) *D. melanogaster sex* DEG expression in pupae (above x-axis) *versus* adults (below x-axis). Clusters in which the sex DEG is expressed without significant sex bias are shown in gray, and clusters with sex-biased expression are shown in shades of orange (female-biased) or purple (male-biased) based on their Iog2-transformed ratio of aggregated female to male gene expression (log2FC). **(F)** Area-proportional Euler diagram showing the numbers of sex DE genes (DEGs) in pupae and adults. **(G)** Sex DEGs that are shared between developmental stages at log2FC>2, showing the number ofclusters with female- or male-bias at each developmental stage for each gene. Columns are clustered hierarchically based on Euclidean distances. The same clustering is also observed for sex DEGs shared between developmental stages at log2FC>1 (not shown).

The extent of sex-biased gene expression further reflected potential transcriptomic convergence between sexes in adults. Under our stringent sex DEG definition and testing on a common gene list, we found more sex DE cluster-gene combinations in pupae than in adults (Figure 5C). As pupae developed into adults, most of these sex DEGs retained a similar expression breadth (Figure 5D-E) but lost their sex-biased expression: while the vast majority of adult sex DEGs were also DE in pupae, most pupae sex DEGs were stage-specific (Figure 5F). Furthermore, for the sex DEGs found in both pupae and adults, the direction of sex-biases show similarity between developmental stages (Figure 5G). Taken together, the complementary datasets suggested that transcriptomic sexual dimorphism in *fru* neurons may decrease during adulthood, as sexes converge to a more similar transcriptome.

## Discussion

Across animals, sexual circuits face a common conundrum: the two sexes share many circuit elements derived from common developmental origins, but they often have drastically different usage and fitness optima of these shared elements. Here, we address how these two opposing forces together shape the extent, distribution, and evolution of sex-biased gene expression within sexual circuits with cellular resolution in the *Drosophila* nervous system. Our sex and species comparisons converge to show that the transcriptomic evolution of sexual circuits occurs primarily through rapid turnover of limited and highly localized sex-biased gene expression among pervasive transcriptomic coupling between sexes.

The limited sex DEGs and extensive transcriptomic coupling between sexes in the sexual circuit defies the *a priori* expectation of widespread sexual dimorphism in gene expression and its evolution. While some earlier bulk RNA-seq studies have hinted at the scarcity of sex-biased gene expression and samples’ lack of a clear transcriptomic separation by sex in the brains across a wide range of species (Goldman and Arbeitman 2007; Catalán et al. 2012; Werling et al. 2016; Naqvi et al. 2019; Brovkina et al. 2021; Rodríguez-Montes et al. 2023; Xie et al. 2024), we extend these findings to a much finer cellular level, and in the parts of the *Drosophila* CNS that orchestrate the most sexually dimorphic behaviors.

The cellular-level analyses provide further clues to this paradoxically limited transcriptomic footprint of sexual conflict. Consistent with the adaptive need for highly cell-type-specific dimorphic gene expression, both sex DEGs and sex-specific species DEGs are typically expressed broadly across many cell clusters but differentially expressed in only a very small subset. This cell-type-specific, sex-specific adaptive need is, however, impeded by genetic constraints that impose shared gene regulatory mechanisms across cell types. Our data illustrates this in two ways: firstly, genes that exhibited sex-biased expression in multiple clusters were highly consistent in the direction of sex bias (Figure 2D, Figure S4B-C), pointing to shared gene regulation of sex-biased gene expression among cell types. Furthermore, the degree of transcriptomic coupling of species DEG between sexes increased with their expression breadth, with species DEGs that evolved in a more sex-specific manner having narrower expression breadths (Figure 4B). This suggests that pleiotropic gene expression imposes limits on the decoupling of gene expression regulation between sexes in a cell-type-specific manner. Given the genetic constraints on sex-specific gene expression adaptation, the nervous system may continue to experience unresolved sexual conflict as new un-decoupleable regulatory changes arise.

In line with the genetic constraint on sex-specific gene expression, the conserved sex DEGs we identified often reflected sex differences in the abundance of fine-grained cell types instead of dimorphic gene expression within homologous cell types (Figure 2H). Clusters with known sex differences in cell types showed more sex DEGs, which might also result from sex differences in cellular composition. A recent scRNA-seq analysis of adult *D. melanogaster* brains also finds evidence for sexual dimorphism via the differential survival of cell types between sexes, rather than substantial transcriptomic remodeling (Allen et al. 2026). Furthermore, recent EM connectome comparisons of neuronal morphology and connectivity in *D. melanogaster* brain defined more sex-specific cell types than sexually dimorphic cell types (Berg et al. 2025; Stürner et al. 2025). These results potentially suggest sex-specific fine-grained cell types as a major resolution of strong sexual conflict, bypassing the potentially unattainable need of extensive cell-type-specific, sex-biased gene expression in shared cell types.

Gonads are highly sexually dimorphic in gene expression, development, anatomy, or cell types across animals (Malone et al. 2006; Small et al. 2009; Mank et al. 2010; Mank and Rideout 2021; Xie et al. 2025). Compared with the brain, some somatic tissues also exhibit a much higher degree of sex-biased gene expression (e.g. Rodríguez-Montes et al. 2023; Sylvestre et al. 2025; Xie et al. 2025). The lack of pervasive transcriptomic dimorphism we see in the part of the nervous system that generates highly sex-specific phenotypic outputs raises interesting features of neural circuits in their response to intralocus sexual conflict. The nervous system is exceptionally heterogenous in cell types. While behaviors typically involve complex and concerted activities of many circuit elements from sensory to motor, sex differences at highly specific circuit nodes may be sufficient in organizing sex-specific sensorimotor transformations and orchestrating sex-specific behavioral outcomes. Furthermore, existing sexual dimorphism, such as sex differences in circuit connectivity, might effectively restrict the behavioral outcome of sex-shared gene expression changes to one sex, shielding the other sex from negative fitness consequences. Consistent with this idea, latent sexual circuitry, where sexual behaviors typical of one sex can be elicited in the other sex under specific conditions, has been reported across a wide range of species and appears to be a common feature of sexual circuits across animals (Kimchi et al. 2007; Clyne and Miesenböck 2008; Rezával et al. 2016; Wei et al. 2018; Michael et al. 2020; Li et al. 2024). Further research may reveal whether the latent circuit also encodes highly species-specific features of sexual behavior, which would be indicative of sex-coupled circuit evolution driving behavioral adaptation in one sex and the capacity to harbor unexpressed or highly context-restricted behavior in the other sex. Finally, sex-shared circuit nodes that are integrated into distinct circuits in each sex and orchestrate sex-specific behavioral outputs may nonetheless experience similar selective pressures and thus share fitness optima. For instance, in *D. melanogaster*, a population of high-order brain neurons are tuned to acoustic features of conspecific courtship song in both males and females, suggesting sex-coupled evolution of auditory perception and processing, but route the acoustic perception to different downstream circuits to elicit sex-specific behavioral responses (Deutsch et al. 2023).

On the gene level, our results provide insights into the molecular programs constrained and poised for sex-specific adaptation in gene expression. Most strikingly, we observed a strong overrepresentation of shared DEGs across sex, species, and FruM-null comparisons, pinpointing a common set of genes, such as GPCR pathway genes (Figure 4F), whose expression is highly flexible across different biological contexts. Moreover, signaling receptor genes, including GPCRs, are enriched among species-specific sex DEGs, whereas transcription factors are enriched among conserved sex DEGs (Figure 2K). This pattern reveals heterogeneous evolutionary dynamics within sex-biased gene expression: the transcription factor code that delineates sex-specific cellular identity is conserved, while effectors, particularly signaling receptor genes, exhibit rapid evolutionary turnover in expression. Consistent with this observation, signaling receptor genes are also enriched in sex-specific species DEGs, in contrast to the transporter and channel activity genes enriched in sex-shared species DEGs (Figure 4C). We hypothesize that the regulatory landscapes of signaling receptor genes, possibly shaped by their context-dependent neural functions (Marti-Solano 2023), make them particularly poised for sex-specific adaptation; whereas other gene classes, such as transporter and channel genes, may experience less intralocus sexual conflict or stronger genetic constraints, which disincentivizes or limits the transcriptional decoupling between sexes, respectively. Emerging scRNA-seq studies in nervous systems of nematodes, fruit flies, and mice have repeatedly identified neuropeptide signaling genes in sex or species comparisons (Chen et al. 2025; Haque et al. 2025; Toker et al. 2025; Walsh et al. 2025), suggesting that sex-specific gene expression evolution of neural signaling genes may be a general mechanism for resolving past and ongoing sexual conflict across organisms.

Finally, our results reveal a possible transcriptomic convergence between sexes in adulthood and highlights developmental timing as an important context to consider sexual dimorphism and the action of sexual conflict (Mank 2017; Tosto et al. 2023). Studies of sex-biased gene expression often focus on the adult stage because phenotypic sex differences are most noticeable at this stage (Mank 2017). However, we found more pronounced transcriptomic sex differences at the mid-pupal stage, a time when FruM expression peaks and sexual circuits have largely completed construction (Lee et al. 2000; Palmateer et al. 2023), than in the adult stage, a time when sex-specific behaviors are being expressed. This transcriptomic convergence towards a highly similar adult state between the sexes mirrors the transcriptomic convergence among neuronal cell types during development seen in the *Drosophila* visual and olfactory systems (Li et al. 2017; Özel et al. 2021). Our results contribute to a growing appreciation of the dynamic nature of the gene regulatory program that shapes neuronal diversity not only among cell types but also between sexes (Kurmangaliyev et al. 2020; Jain et al. 2022; Rodríguez-Montes et al. 2023; Cachero et al. 2025).

Although our study provides the finest cellular resolution for sex comparisons of transcriptomic evolution in any system to date, limitations and open questions remain. Given the highly localized nature of sexual dimorphism even within *fru* neurons, our current cellular coverage is modest. Increased coverage could uncover more localized sex DEGs, such as those present in only one or a few neurons. Nonetheless, we expect that the relative abundance of sex DEGs, species DEGs, and FruM DEGs robust to coverage. Furthermore, the observed temporal dynamics of sexual dimorphism raises the possibility that the degree of transcriptomic decoupling between sexes and the molecular programs providing sex-specific evolvability in earlier circuit development may differ from those in the adult stage. For example, cell-surface molecules play key roles in sexually dimorphic, cell-type-specific neuronal wiring (Li et al. 2020; Kim and Kim 2022; Lyu et al. 2026). Their expressions are highly cell-type–specific and temporally dynamic during neuronal development (Kalish et al. 2018; Lo Giudice et al. 2019; Jain et al. 2022; Choi et al. 2023). Although we did not detect significant enrichment of these genes among pupal sex DEGs, they may nevertheless be a major contributor of sex-specific, cell-type-specific adaptation in neuronal wiring in a transient manner that is not fully captured by a single developmental snapshot. Finally, while we address sexual dimorphism at the level of the transcriptome in this study, how these gene expression changes are translated into sex-shared or sex-specific outcomes at the levels of neurons, circuits, or behaviors remain unknown. Exciting opportunities lie in future integration of comparisons across various modalities such as neuronal morphology and circuit connectivity (Deutsch et al. 2025; Stürner et al. 2025) to infer the organization, evolution, and behavioral consequences of sexual circuits under sex-shared developmental origin and intralocus sexual conflict across biological levels.

## Methods

### Fly stocks and husbandry

Flies were maintained on cornmeal-agar-yeast medium (Fly Food B, Bloomington Recipe, Lab Express) at 23°C and 50% humidity on a 12 hour light and dark cycle.

### Generation of genetic reagents

*D. melanogaster* and *D. yakuba fru-GAL4* were generated through CRISPR/Cas9-mediated knock-in using the same methods as in (Ye et al. 2024). For each species, Gal4 was inserted in-frame at the endogenous start codon of *fruM*, with a 274 bp deletion starting from ATG generated. The *fru-GAL4* alleles are expected to function as FruM-null, which was confirmed by the FruM-null behavioral phenotypes (i.e., male-male courtship (Ryner et al. 1996; Demir and Dickson 2005)) in both species and anti-FruM immunostaining in *D. melanogaster* (Figure S1B).

### Sample collection and sequencing

D. melanogaster control males and females had the genotype UAS-myrGFP/+; fru-GAL4 Mhc-DsRed/UAS-nls-tdTomato. D. melanogaster FruM-null males had the genotype UAS-myrGFP/+; fru-GAL4 Mhc-DsRed UAS-nls-tdTomato/fru-GAL4 Mhc-DsRed. D. yakuba control males and females had the genotype fru-GAL4 Mhc-DsRed/UAS-nls-tdTomato. Flies were collected within 16 hours of eclosion and housed in single-sex vials of 10-15 flies per vial until dissection. All flies were 4-6 days old at the time of dissection.

Our dissection, cell dissociation, FACS sorting and scRNA-seq library preparation protocol was previously described in (Walsh et al. 2025). Briefly, whole CNS of 40-75 ice-anesthetized flies were dissected in ice-cold Schneider’s medium (Gibco, 21720024) within 1 hour of anesthesia. CNS tissue were then transferred to 500 µl of dispase (1 mg/ml Liberase DH (Roche, 5401054001) in adult hemolymph saline (AHS) buffer) and digested at room temperature for 50 min on a nutator, before dispase was removed and digested tissue was washed 3 times with ice-cold Schneider’s medium. For cell dissociation, we added 500 µl of ice-cold 1X phosphate buffered saline (PBS; Thermo Fisher) with 2% bovine serum albumin (BSA; Thermo Fisher, B14) to the tissue, and first used a P200 pipette tip (coated with fetal bovine serum (FBS; Fisher Scientific, SH3007103) to prevent cells from sticking) to pipet the sample up and down 100 times, then used a 3 ml syringe with 25 gauge needle (coated with FBS) to draw the sample up and down another 20 times. The cell suspension was then transferred into a low-bind culture tube and FACS-sorted for nls-tdTomato+ (*fru*+) cells using an Aria Cell Sorter with 100 µm nozzle and 1.25 drop precision setting. A FACS negative control sample from flies without nls-tdTomato-labelled *fru* neurons were always prepared alongside sequencing samples to establish baseline fluorescence level and confirm the threshold for tdTomato+ cells.

#### scRNA-seq library preparation and sequencing

After FACS sorting, samples were gently centrifuged at 300 g for 3 min at 4°C, and excess supernatant was removed to reach a final concentration of 1000 cells/µl. Samples were then submitted to the Center for Applied Genomics at the Children’s Hospital of Philadelphia’s Research Institute for library preparation using the 10X Genomics Single Cell 3’ Library and Gel Bead Kit v3.1 following the manufacturer’s instructions, and sequenced on Illumina NovaSeq 6000. Two replicates were prepared for each sample type. With the exception of one *D. yakuba* female and one *D. yakuba* male sample, all other samples were prepared on separate days.

#### Bulk RNA-seq library preparation and sequencing

Three replicates each were prepared for *D. melanogaster* males and *D. yakuba* males over 2 days, with each replicate deriving from a separate cross. Two *D. melanogaster* and one *D. yakuba* sample were prepared in batch 1, and the other samples in batch 2.

After FACS sorting, samples were centrifuged at 0.9 g for 15 min at 4°C. The supernatant was removed and TRIzol was added to the pellet at 1 µl/1000 cells. The samples were then frozen on dry ice and stored at -80°C before all six samples were submitted to Azenta at the same time for RNA-extraction, library preparation and sequencing. SMART-Seq v4 Ultra Low Input Kit for Sequencing was used for full-length cDNA synthesis and amplification (Clontech), and Illumina Nextera XT library was used for sequencing library preparation. Briefly, cDNA was fragmented, and adaptor was added using Transposase, followed by limited-cycle PCR to enrich and add index to the cDNA fragments. The sequencing library was validated on the Agilent TapeStation (Agilent Technologies), and quantified by using Qubit 3.0 Fluorometer (Invitrogen) as well as by quantitative PCR (KAPA Biosystems). The sequencing libraries were multiplexed and clustered onto a flowcell on the Illumina NovaSeq instrument according to manufacturer’s instructions. The samples were sequenced using a 2×150 bp paired end configuration. Image analysis and base calling were conducted by the NovaSeq Control Software. Raw sequence data (.bcl files) generated from Illumina NovaSeq was converted into fastq files and de-multiplexed using Illumina bcl2fastq 2.20 software. One mis-match was allowed for index sequence identification.

### scRNA-seq data processing and analysis

We used CellRanger’s (v7.1.0) “count” function to map reads to the *D. melanogaster* (Assembly GCA_000001215.4) and *D. yakuba* (Assembly GCA_016746365.2) genomes, respectively. Cell expression matrices were processed in Seurat (v5.1.0) (Hao et al. 2024). We removed cells that expressed fewer than 200 (low-quality cells) or more than 4,000 unique genes (potential doublets), more than 20,000 transcripts (potential doublets), or more than 5% mitochondrial counts (low-quality or dying cells). We considered only 1:1 orthologs identified by best reciprocal BLAST hits that were expressed in ≥3 cells in each sample, and identified 11,051 genes across all samples. We then merged all samples (“merge”) and followed the standard Seurat workflow with default settings to normalize (“NormalizeData”) and scale (“ScaleData”, regressing out the percentage of mitochondrial counts and number of transcripts) the data, and performed principal component analysis (PCA; “RunPCA”) to identify the statistical significance of PCs. The PCA was then used to inform anchor-based CCA integration across samples (“IntegrateLayers”), and the integrated data was clustered (“FindNeighbors” using the first 30 PCs and “FindClusters”) and visualized through UMAP dimensionality reduction (“RunUMAP”). We tried clustering resolutions 1, 2, and 3, and found that higher resolutions simply introduced more granularity by subdividing larger clusters into smaller ones, without broad impacts on cluster assignment. Therefore, we chose the intermediate resolution of 2, which produced 102 clusters. Annotation of these clusters (see “Cell and cluster annotation”) revealed that each annotated cluster generally corresponded to one anatomical population or VNC hemilineage, suggesting that the resolution is appropriate for our dataset. Quality control on the distribution of gene counts, transcript counts, mitochondrial counts, and *fru* expression were performed according to standard Seurat workflow. The quality metrics (Figure S1D, Supplementary Table 1) were comparable to or outperformed previous studies of *Drosophila* CNS (Davie et al. 2018; Allen et al. 2020; Palmateer et al. 2023; Lee et al. 2025). Four clusters with <90% of cells expressing any neuronal marker genes (*elav, brp, Syt1,* or *CadN*) or with <50% of cells expressing *fru* were excluded from further analysis.

#### Cell and cluster annotation

CNS region (brain or VNC) and fast-acting neurotransmitter identity were annotated for each cell based on marker gene expression. VNC cells were defined as those expressing any of the segment identity Hox genes *Antp*, *Ubx*, *abd-A*, or *Abd-B*, or a higher expression of the VNC marker *tsh* than the brain marker *oc*. Brain cells were non-VNC cells that expressed any of the neuronal markers *elav*, *brp*, *Syt1* or *CadN*, or *oc*. Fast-acting neurotransmitter identities were annotated with the following markers: cholinergic neurons with *VAChT* or *ChAT*, glutamatergic neurons with *VGlut*, and GABAergic neurons with *Gad1*. Consensus CNS region and fast-acting neurotransmitter annotations were reached if more than 75% of cells in all 4 wildtype samples had the same annotation.

We annotated visual system populations using the 96 hr APF cells from a visual system atlas ((Kurmangaliyev et al. 2020); atlas V1.1; https://zenodo.org/records/8111612). *D. melanogaster* female and male samples were used for comparison with the visual system atlas, with all genes detected in these samples (not limited to 1:1 orthologs). The mean normalized expression values were calculated for each cluster in both datasets. We then identified up to 10 best marker genes for each *fru* cluster that with detection rate>0.25, log2FC>3, and adjusted p<0.01 that was also present in the visual system atlas, and used the log-transformed expression values of these genes to calculate pairwise Pearson’s correlation coefficients between clusters across datasets. We considered matches with Pearson’s r>0.8. All but one matched cluster had reciprocal best matches between datasets, and cluster 40 matched to LLPC1, 2, and 3. Overall, we matched 21 clusters to visual system populations: Dm8, Dm9, Lawf2, LC10a, LLPC1/2/3, LPLC2, T1, T4/T5, Tm3, TmY3, N33, N55, N58, N70, N76, N82, N88, N93, N102, N122, N140. We further verified these matches by their expression of genetic markers reported for each optic lobe cluster (Kurmangaliyev et al. 2020; Naidu et al. 2020; Özel et al. 2021). Cluster 63 (T4/T5) was not included in the analysis because it did not meet the *fru* expression threshold.

Other clusters were annotated with their anatomical populations using markers from the literature. Brain populations aDT-a, aDT-b, aDT-h, aSP-a, aSP-k, pIP-e, pIP-z, pMP-b, pMP-e, pMP-z, and PAM were annotated using on markers from (Allen et al. 2026); clock neurons and ɑβ and ɣ Kenyon Cells using markers from (Palmateer et al. 2023); and VNC hemilineages using markers from (Cachero et al. 2025). Clusters where all marker genes occupied a region in the cluster UMAP space (i.e., the anatomical population constituted a subset of the cluster) were labeled as containing (∋) that anatomical population. Monoaminergic clusters were identified based on the expression of *Vmat*, and further annotated by their monoamine identities with the following markers: dopaminergic with *DAT* or *ple*, serotonergic with *SerT* or *Trhn*, tyraminergic/octopaminergic with *Tdc2* or *Tbh*, and histaminergic with *Hdc*. Cluster 81 was composed of a mixture of serotonergic, tyraminergic/octopaminergic, and histaminergic cells, and was labeled “Other monoaminergic”.

#### Cell type abundance (cluster size)

*D. melanogaster* female and male cluster sizes were normalized to the equivalent of one individual based on empirical counts of *fru* neurons the CNS (excluding the mushroom body and visual system) of *fru^GAL4^ UAS-nlacZ* adult females and males (3,333 and 3,157, respectively; (Stockinger et al. 2005)). In *D. yakuba* and in *D. melanogaster* FruM-null males, we normalized cluster sizes under the assumption that the median cluster size ratio between species within each sex or between FruM-null and wildtype males, respectively, was 1.

As a measure of sexual dimorphism or species divergence in cluster size, we calculated the log2 fold-change of normalized cluster size between sexes and species, respectively. Clusters with fewer than 5 cells in both sexes or species were excluded from this analysis. Statistical significance of cluster size difference was performed using the Python package scCODA with automatic reference cluster selection (v0.1.9) (Büttner et al. 2021). We reported clusters and subclusters with non-0 effect estimates under FDR<0.2 or 0.05 as potential candidates with sex, species, or FruM differences in cluster sizes.

#### Transcriptomic correlation

To compare homologous clusters’ transcriptomic correlation between sexes and species, we aggregated normalized gene expression (“AggregateExpression”) for each sample type and cluster across replicates, selected the lists of genes that were expressed in >10% of the cells in that particular cluster in either sample type, and used the gene list to calculate Kendall’s tau correlation coefficient between sample types. Transcriptomic correlation between replicates were also calculated for each cluster in each condition, using the list of genes that were expressed in >10% of the cells in that cluster and sample type. We found that cluster size (log-transformed), inter-replicate transcriptomic correlation, and, to a lesser extent, number of genes used in Kendall’s tau calculation, had significant confounding effects on transcriptomic correlation. Therefore, we used linear models to either 1) include these variables as covariates when comparing distributions of transcriptomic correlations between sample types; or 2) regress out these variables and use the residuals when calculating correlations between inter-sex or inter-species transcriptomic correlations.

#### Differential gene expression

We used Seurat’s “FindMarkers” function with the MAST method to perform differential gene expression analyses while accounting for mitochondrial content (percent.mt) and number of transcripts (nCount_RNA) as latent variables. We note that MAST uses a hurdle model to model gene expression in scRNA-seq data, and produces log2FC values that are different from mean-based log2FC values. Genes that were expressed in <1% of cells in either species were not tested for differential expression to avoid artifacts due to genome annotation quality discrepancies. We only considered DEGs with adjusted p<0.05, and further filtered them based on log2 fold-change (>0, >1, or >2). In the main text, we focused on stringent DEGs with log2 fold-change>2 unless otherwise specified. Overlap between sets of DE genes or cluster-gene combinations were defined as those with p<0.05 in both sets and log2FC>2 (stringent) or >1 (relaxed) in at least one set in order to mitigate underestimation arising from threshold sensitivity. Significance of overlaps between sets of genes or cluster-gene combinations were tested using the hypergeometric test. In Figure 2H and Figure S4E, the large number of sex DEGs detected in cluster 0 (ɣ Kenyon cells) might be attributable to the higher statistical power of a larger cluster of potentially highly homogenous cell types.

#### GO enrichment

The R packages clusterProfiler (v4.10.1; (Wu et al. 2021)) and org.Dm.eg.db (v3.18.0) were used to perform GO enrichment analysis with user-defined genes of interest and gene universe. HTSanalyzeR’s (v2.3.5; (Wang et al. 2011)) drosoAnnotationConvertor function was used to convert gene names to Entrez IDs.

#### Re-analysis of pupal data

Reads from Palmateer et al. (Palmateer et al. 2023) were downloaded from NCBI Gene Expression Omnibus (ID GSE160370), processed, and analyzed using the same methods as our adult data. We examined the expression of previously identified maturation transcription factors *Hr3* and *Hr4* (Jain et al. 2022; Elkahlah et al. 2025) and 11 additional genes with dynamic and pan-neuronally coordinated expression during pupal development: *Blimp-1, Hr38, elav, Rbp9, bol, Snap25, brp, Rbp, nSyb, Syt12*, and *Drep2* (Kurmangaliyev et al. 2020). Sex-biased expression of these 13 genes were minimal, with only 18 cluster-gene combinations involving these genes having an adjusted p-value<0.05, and 1 further meeting log2FC>1 (Figure S10). Therefore, we did not observe any clear neuron maturation differences between male and female pupal data that might confound the detection of sex-biased gene expression.

The pupal and wildtype adult *D. melanogaster* datasets integrated poorly: UMAP of the integrated dataset showed clear separation by developmental stages, and transcriptomic similarity between pupal and adult clusters did not reveal clear one-to-one relationships that would facilitate identification of homologous clusters across developmental stages. Additionally, label transfer (“FindTransferAnchors” and “MapQuery”) using wildtype adult *D. melanogaster* datasets as the reference and pupal datasets as the query yielded only 35 (out of 98) clusters with reciprocal best matches and generally low mapping scores (0.654±0.232). These results might stem from dramatic transcriptomic reprogramming of *fru* neurons during development, and be further contributed by potential differences in the *fru-GAL4* reagents and/ or library preparation chemistries and techniques. Consequently, both datasets were analyzed separately without integration. At resolution=2, our re-analysis of the pupal data contained 75 clusters, three of which were excluded because <50% of the cells expressed *fru* and/or <90% of the cells expressed any neuronal marker (*elav*, *nSyb, brp, Syt1*, or *CadN*).

### Bulk RNA-seq data processing and analysis

Reads were trimmed of their adapters using trimmomatic (v0.39) with the following settings: ILLUMINACLIP:NexteraPE-PE.fa:2:30:10:2:True LEADING:3 TRAILING:3 MINLEN:36, mapped to their respective genomes (*D. melanogaster a*ssembly GCA_000001215.4 and *D. yakuba* assembly GCA_016746365.2) using STAR with default settings, and sorted and indexed using samtools with default settings. Mapped reads were then assigned to exons and counted using featureCounts with the “-largestOverlap” and “-p --countReadPairs” options. 18.9-24.9M reads were assigned for each sample, representing a 67.7-76.5% read assignment rate.

We used the standard DESeq2 (v1.42.1; (Love et al. 2014)) workflow to identify species DEGs among the 12,467 1:1 orthologs that were expressed in this dataset, while accounting for batch effect. Log2 fold-changes were shrunk using the apeglm method (v1.24.0; (Zhu et al. 2019)).

### General data analysis and statistics

Data analysis was performed in R (v4.3.2) with the following packages: tidyverse (v2.0.0; (Wickham et al. 2019)), lme4 (v1.1.37 ;(Bates et al. 2015)), lmerTest (v3.1.3 ;(Kuznetsova et al. 2017)), emmeans (v1.11.0 ; https://CRAN.R-project.org/package=emmeans), clustree (v0.5.1; (Zappia and Oshlack 2018)), ape (v5.8.1; (Paradis and Schliep 2019)), and phytools (v2.4.4; (Revell 2024)). Linear models (or linear mixed models where appropriate) were fitted to the data and the statistical significance of predictors were assessed with two-sided ANOVA either with post hoc Tukey test or p-values were Bonferroni-corrected. When there was significant deviation from the assumptions of linear models, the non-parametric Spearman correlation and Wilcoxon rank-sum test was used.

### Immunostaining

In Figure S1A, whole CNSs were dissected in 1X PBS within 60 min of ice anesthesia. Samples were then fixed in 4% paraformaldehyde (PFA) for 35 min, washed 3 times in PBS with 1% Triton X-100 (PBTX), and blocked in PBTX with 5% normal goat serum (NGS) for 1.5 hr at room temperature, before incubating with primary antibodies diluted in PBTX with 5% NGS at 4°C overnight. Samples were then washed 3 times in PBTX for 30 min each wash, and incubated with secondary antibodies diluted in PBTX with 5% NGS at 4°C overnight. Lastly, after 3 washes in PBTX for 30 min each, the samples were mounted with ProLong™ Gold Antifade Mountant (P10144, Invitrogen) on poly-L-lysine coated coverslips and sealed on all sides with nail polish. The primary antibodies used were chicken-anti-GFP (1:600, ab13970, Abcam) and mouse-anti-nc82 (1:30, DHSB). The secondary antibodies used were goat-anti-chicken/AF488 (1:500, A-11039, Thermo Fisher) and goat-anti-mouse/AF647 (1:500, A-28181, Thermo Fisher). In Figure S1B, whole CNSs were immunostained as described in (Coleman et al. 2024) after dissection in 1X PBS and mounted with ProLong™ Gold Antifade Mountant (P10144, Invitrogen). In addition to the primary antibodies used in Figure S1A, the samples were also incubated with the primary antibody rabbit-anti-FruM (1:1000, gift from Vanessa Ruta and Rory Coleman); the secondary antibodies used were goat-anti-chicken/AF488 (1:500, A-11039, Thermo Fisher), goat-anti-mouse/AF568 (1:500, A-11031, Thermo Fisher) and goat-anti-rabbit/ AF647 (1:500, PIA32733TR, Invitrogen). Confocal images were taken on a Leica DMi8 microscope with a TCS SP8 Confocal system at 40x, and processed with FIJI (v2.16.0/1.54p).

## Data availability

The dataset generated here will be available upon publication.

## Acknowledgments

We thank Sarah Flanagan, Junhyong Kim, Li Zhao, Troy Shirangi, Erol Akçay, and Chris Large for helpful discussions, and lab members for CNS dissections and discussions. We are grateful to Rory Coleman and Vanessa Ruta for the antibody reagent. This project was supported by the NIH grant R35GM142678 to Y.D.

## Author contributions

D.S.C and Y.D. conceived the study. Y.D. generated genetic reagents. D.S.C. collected and analyzed sequencing data. H.G. collected imaging data. Y.Z.K. annotated optic lobe clusters. D.S.C and Y.D. wrote the manuscript with contributions from all authors.

## References

Allen AM, Neville MC, Birtles S, Croset V, Treiber CD, Waddell S, Goodwin SF. 2020. A single-cell transcriptomic atlas of the adult ventral nerve cord. Elife. 9. doi:10.7554/eLife.54074. http:// dx.doi.org/10.7554/eLife.54074.

Allen AM, Neville MC, Nojima T, Alejevski F, Agarwal D, Sims D, Goodwin SF. 2025 Dec 19. A high-resolution atlas of the brain predicts lineage and birth order underlying neuronal identity. Cell Genom.:101103.

Allen AM, Neville MC, Nojima T, Alejevski F, Goodwin SF. 2026 Jan 12. Differential neuronal survival defines a novel axis of sexual dimorphism in the Drosophila brain. Cell Genom.:101125.

Anand A, Villella A, Ryner LC, Carlo T, Goodwin SF, Song HJ, Gailey DA, Morales A, Hall JC, Baker BS, et al. 2001. Molecular genetic dissection of the sex-specific and vital functions of the Drosophila melanogaster sex determination gene fruitless. Genetics. 158(4):1569–1595.

Aranha MM, Vasconcelos ML. 2018. Deciphering Drosophila female innate behaviors. Curr Opin Neurobiol. 52:139–148.

Arnqvist G, Rowe L. 2013. Sexual Conflict. Princeton University Press.

Asahina K. 2018. Sex differences in behavior: Qualitative and Quantitative Dimorphism. Curr Opin Physiol. 6:35–45.

Ayroles JF, Carbone MA, Stone EA, Jordan KW, Lyman RF, Magwire MM, Rollmann SM, Duncan LH, Lawrence F, Anholt RRH, et al. 2009. Systems genetics of complex traits in Drosophila melanogaster. Nature Genetics. 41(3):299–307.

Baker CA, Guan X-J, Choi M, Murthy M. 2024. The role of in specifying courtship behaviors across divergent species. Sci Adv. 10(11):eadk1273.

Bates D, Mächler M, Bolker B, Walker S. 2015. Fitting linear mixed-effects models Usinglme4. J Stat Softw. 67(1). doi:10.18637/jss.v067.i01. http://www.jstatsoft.org/v67/i01/.

Bedhomme S, Chippindale AK. 2007. Irreconcilable differences: when sexual dimorphism fails to resolve sexual conflict. In: Sex, Size and Gender Roles. Oxford University PressOxford. p. 185–194.

Berg S, Beckett IR, Costa M, Schlegel P, Januszewski M, Marin EC, Nern A, Preibisch S, Qiu W, Takemura S-Y, et al. 2025. Sexual dimorphism in the complete connectome of the male central nervous system. bioRxiv. doi:10.1101/2025.10.09.680999. http://dx.doi.org/10.1101/2025.10.09.680999.

Bertossa RC, van de Zande L, Beukeboom LW. 2009. The Fruitless gene in Nasonia displays complex sex-specific splicing and contains new zinc finger domains. Mol Biol Evol. 26(7):1557–1569.

Billeter J-C, Rideout EJ, Dornan AJ, Goodwin SF. 2006. Control of male sexual behavior in Drosophila by the sex determination pathway. Curr Biol. 16(17):R766–76.

Biswas P, Rideout EJ. 2025. Sex-biased expression of enteroendocrine cell-derived hormones contributes to higher fat storage in Drosophila females. eLife. doi:10.7554/elife.109426.1. https://elifesciences.org/reviewed-preprints/109426v1.

Bonduriansky R, Chenoweth SF. 2009. Intralocus sexual conflict. Trends Ecol Evol. 24(5):280–288.

Bontonou G, Saint-Leandre B, Kafle T, Baticle T, Hassan A, Sánchez-Alcañiz JA, Arguello JR. 2024. Evolution of chemosensory tissues and cells across ecologically diverse Drosophilids. Nat Commun. 15(1):1047.

Brovkina MV, Duffié R, Burtis AEC, Clowney EJ. 2021. Fruitless decommissions regulatory elements to implement cell-type-specific neuronal masculinization. PLoS Genet. 17(2):e1009338.

Büttner M, Ostner J, Müller CL, Theis FJ, Schubert B. 2021. scCODA is a Bayesian model for compositional single-cell data analysis. Nat Commun. 12(1):6876.

Cachero S, Mitletton M, Beckett IR, Marin EC, Serratosa Capdevila L, Gkantia M, Lacin H, Jefferis GSXE, Donà E. 2025. A developmental atlas of the *Drosophila* nerve cord uncovers a global temporal code for neuronal identity. bioRxiv. doi:10.1101/2025.07.16.664682. http://biorxiv.org/lookup/doi/10.1101/2025.07.16.664682.

Cachero S, Ostrovsky AD, Yu JY, Dickson BJ, Jefferis GSXE. 2010. Sexual dimorphism in the fly brain. Curr Biol. 20(18):1589–1601.

Catalán A, Hutter S, Parsch J. 2012. Population and sex differences in Drosophila melanogaster brain gene expression. BMC Genomics. 13:654.

Catalán A, Macias-Muñoz A, Briscoe AD. 2018. Evolution of Sex-Biased Gene Expression and Dosage Compensation in the Eye and Brain of Heliconius Butterflies. Mol Biol Evol. 35(9):2120–2134.

Chen J, Jin S, Chen D, Cao J, Ji X, Peng Q, Pan Y. 2021. tunes functional flexibility of courtship circuitry during development. Elife. 10. doi:10.7554/eLife.59224. http://dx.doi.org/10.7554/ eLife.59224.

Chen J, Richardson PR, Kirby C, Eddy SR, Hoekstra HE. 2025. Cellular evolution of the hypothalamic preoptic area of behaviorally divergent deer mice. Elife. 13. doi:10.7554/eLife.103109. http://dx.doi.org/10.7554/eLife.103109.

Choi JS, Ayupe AC, Beckedorff F, Catanuto P, McCartan R, Levay K, Park KK. 2023. Single-nucleus RNA sequencing of developing superior colliculus identifies neuronal diversity and candidate mediators of circuit assembly. Cell Rep. 42(9):113037.

Clyne JD, Miesenböck G. 2008. Sex-specific control and tuning of the pattern generator for courtship song in Drosophila. Cell. 133(2):354–363.

Coleman RT, Morantte I, Koreman GT, Cheng ML, Ding Y, Ruta V. 2024. A modular circuit coordinates the diversification of courtship strategies. Nature. 635(8037):142–150.

Cooke B, Hegstrom CD, Villeneuve LS, Breedlove SM. 1998. Sexual differentiation of the vertebrate brain: principles and mechanisms. Front Neuroendocrinol. 19(4):323–362.

Davie K, Janssens J, Koldere D, De Waegeneer M, Pech U, Kreft Ł, Aibar S, Makhzami S, Christiaens V, Bravo González-Blas C, et al. 2018. A Single-Cell Transcriptome Atlas of the Aging Drosophila Brain. Cell. 174(4):982–998.e20.

Demir E, Dickson BJ. 2005. fruitless splicing specifies male courtship behavior in Drosophila. Cell. 121(5):785–794.

Deutsch D, Clemens J, Thiberge SY, Guan G, Murthy M. 2023. Shared Song Detector Neurons in Drosophila Male and Female Brains Drive Sex-Specific Behaviors. Curr Biol. 33(17):3796–3800.

Deutsch D, Matsliah A, Wang K, Dorkenwald S, Mondal A, Burke A, Hebditch J, Gager J, Yu S-C, Sterling A, et al. 2025. Sexually-dimorphic neurons in the whole-brain connectome. bioRxiv. doi:10.1101/2025.06.10.658788. http://dx.doi.org/10.1101/2025.06.10.658788.

Deutsch D, Pacheco D, Encarnacion-Rivera L, Pereira T, Fathy R, Clemens J, Girardin C, Calhoun A, Ireland E, Burke A, et al. 2020. The neural basis for a persistent internal state in females. Elife. 9. doi:10.7554/eLife.59502. http://dx.doi.org/10.7554/eLife.59502.

Ding Y, Lillvis JL, Cande J, Berman GJ, Arthur BJ, Long X, Xu M, Dickson BJ, Stern DL. 2019. Neural Evolution of Context-Dependent Fly Song. Curr Biol. 29(7):1089–1099.e7.

van Doorn GS. 2009. Intralocus sexual conflict. Ann N Y Acad Sci. 1168:52–71.

Elkahlah N, Lin Y, Shirangi TR, Clowney EJ. 2025. Hierarchical diversification of instinctual behavior neurons by lineage, birth order, and sex. bioRxiv. doi:10.1101/2025.06.03.657692. http://dx.doi.org/10.1101/2025.06.03.657692.

Ellegren H, Parsch J. 2007. The evolution of sex-biased genes and sex-biased gene expression. Nat Rev Genet. 8(9):689–698.

Freeman MR. 2015. Drosophila Central Nervous System Glia. Cold Spring Harb Perspect Biol. 7(11). doi:10.1101/cshperspect.a020552. http://dx.doi.org/10.1101/cshperspect.a020552.

Gailey DA, Billeter J-C, Liu JH, Bauzon F, Allendorfer JB, Goodwin SF. 2006. Functional conservation of the fruitless male sex-determination gene across 250 Myr of insect evolution. Mol Biol Evol. 23(3):633–643.

Goldman TD, Arbeitman MN. 2007. Genomic and functional studies of Drosophila sex hierarchy regulated gene expression in adult head and nervous system tissues. PLoS Genet. 3(11):e216.

Hao Y, Stuart T, Kowalski MH, Choudhary S, Hoffman P, Hartman A, Srivastava A, Molla G, Madad S, Fernandez-Granda C, et al. 2024. Dictionary learning for integrative, multimodal and scalable single-cell analysis. Nat Biotechnol. 42(2):293–304.

Haque R, Setty H, Lorenzo R, Stelzer G, Rotkopf R, Salzberg Y, Goldman G, Kumar S, Halber SN, Leifer AM, et al. 2025. Decoding sexual dimorphism of the sex-shared nervous system at single-neuron resolution. bioRxiv. doi:10.1101/2024.12.27.630541. http://dx.doi.org/10.1101/2024.12.27.630541.

Harrison PW, Wright AE, Zimmer F, Dean R, Montgomery SH, Pointer MA, Mank JE. 2015. Sexual selection drives evolution and rapid turnover of male gene expression. Proc Natl Acad Sci U S A. 112(14):4393–4398.

Hoopfer ED, Jung Y, Inagaki HK, Rubin GM, Anderson DJ. 2015. P1 interneurons promote a persistent internal state that enhances inter-male aggression in Drosophila. Elife. 4. doi:10.7554/eLife.11346. http://dx.doi.org/10.7554/eLife.11346.

Innocenti P, Morrow EH. 2010. The sexually antagonistic genes of Drosophila melanogaster. PLoS Biol. 8(3):e1000335.

Ito H, Fujitani K, Usui K, Shimizu-Nishikawa K, Tanaka S, Yamamoto D. 1996. Sexual orientation in Drosophila is altered by the satori mutation in the sex-determination gene fruitless that encodes a zinc finger protein with a BTB domain. Proc Natl Acad Sci U S A. 93(18):9687–9692.

Jain S, Lin Y, Kurmangaliyev YZ, Valdes-Aleman J, LoCascio SA, Mirshahidi P, Parrington B, Zipursky SL. 2022. A global timing mechanism regulates cell-type-specific wiring programmes. Nature. 603(7899):112–118.

Jiang X, Ma M, Sun M, Chen J, Pan Y. 2023. Molecular and neuronal mechanisms governing sexually dimorphic prioritization of innate behaviors. bioRxiv. doi:10.1101/2023.12.04.569869. http://biorxiv.org/lookup/doi/10.1101/2023.12.04.569869.

Jiang X, Sun M, Chen J, Pan Y. 2024. Sex-Specific and State-Dependent Neuromodulation Regulates Male and Female Locomotion and Sexual Behaviors. Research (Wash D C). 7:0321.

Kalish BT, Cheadle L, Hrvatin S, Nagy MA, Rivera S, Crow M, Gillis J, Kirchner R, Greenberg ME. 2018. Single-cell transcriptomics of the developing lateral geniculate nucleus reveals insights into circuit assembly and refinement. Proc Natl Acad Sci U S A. 115(5):E1051–E1060.

Khodursky S, Svetec N, Durkin SM, Zhao L. 2020. The evolution of sex-biased gene expression in the brain. Genome Res. 30(6):874–884.

Kimchi T, Xu J, Dulac C. 2007. A functional circuit underlying male sexual behaviour in the female mouse brain. Nature. 448(7157):1009–1014.

Kim D-H, Jang Y-H, Yun M, Lee K-M, Kim Y-J. 2024. Long-term neuropeptide modulation of female sexual drive via the TRP channel in. Proc Natl Acad Sci U S A. 121(10):e2310841121.

Kim D, Kim B. 2022. Anatomical and Functional Differences in the Sex-Shared Neurons of the Nematode. Front Neuroanat. 16:906090.

Kimura K-I, Hachiya T, Koganezawa M, Tazawa T, Yamamoto D. 2008. Fruitless and doublesex coordinate to generate male-specific neurons that can initiate courtship. Neuron. 59(5):759–769.

Kimura K-I, Ote M, Tazawa T, Yamamoto D. 2005. Fruitless specifies sexually dimorphic neural circuitry in the Drosophila brain. Nature. 438(7065):229–233.

Kimura K-I, Sato C, Koganezawa M, Yamamoto D. 2015. Drosophila ovipositor extension in mating behavior and egg deposition involves distinct sets of brain interneurons. PLoS One. 10(5):e0126445.

Kim WJ, Jan LY, Jan YN. 2013. A PDF/NPF neuropeptide signaling circuitry of male Drosophila melanogaster controls rival-induced prolonged mating. Neuron. 80(5):1190–1205.

Kitamoto T. 2002. Conditional disruption of synaptic transmission induces male-male courtship behavior in Drosophila. Proc Natl Acad Sci U S A. 99(20):13232–13237.

Knoedler JR, Inoue S, Bayless DW, Yang T, Tantry A, Davis C-H, Leung NY, Parthasarathy S, Wang G, Alvarado M, et al. 2024. A functional cellular framework for sex and estrous cycle-dependent gene expression and behavior. Cell. 187(14):3781.

Koganezawa M, Kimura K-I, Yamamoto D. 2016. The Neural Circuitry that Functions as a Switch for Courtship versus Aggression in Drosophila Males. Curr Biol. 26(11):1395–1403.

Kopp A, Duncan I, Godt D, Carroll SB. 2000. Genetic control and evolution of sexually dimorphic characters in Drosophila. Nature. 408(6812):553–559.

Kurmangaliyev YZ, Yoo J, Valdes-Aleman J, Sanfilippo P, Zipursky SL. 2020. Transcriptional Programs of Circuit Assembly in the Drosophila Visual System. Neuron. 108(6):1045–1057.e6.

Kuznetsova A, Brockhoff PB, Christensen RHB. 2017. LmerTest package: Tests in linear mixed effects models. J Stat Softw. 82(13). doi:10.18637/jss.v082.i13. http://www.jstatsoft.org/v82/i13/.

Lande R. 1980. SEXUAL DIMORPHISM, SEXUAL SELECTION, AND ADAPTATION IN POLYGENIC CHARACTERS. Evolution. 34(2):292–305.

Lee D, Shahandeh MP, Abuin L, Benton R. 2025. Comparative single-cell transcriptomic atlases of drosophilid brains suggest glial evolution during ecological adaptation. PLoS Biol. 23(4):e3003120.

Lee G, Foss M, Goodwin SF, Carlo T, Taylor BJ, Hall JC. 2000. Spatial, temporal, and sexually dimorphic expression patterns of the fruitless gene in the Drosophila central nervous system. J Neurobiol. 43(4):404–426.

Li C-Y, Bowers JM, Alexander TA, Behrens KA, Jackson P, Amini CJ, Juntti SA. 2024. A pheromone receptor in cichlid fish mediates attraction to females but inhibits male parental care. Curr Biol. 34(17):3866–3880.e7.

Li H, Horns F, Wu B, Xie Q, Li J, Li T, Luginbuhl DJ, Quake SR, Luo L. 2017. Classifying Drosophila Olfactory Projection Neuron Subtypes by Single-Cell RNA Sequencing. Cell. 171(5):1206–1220.e22.

Li J, Han S, Li H, Udeshi ND, Svinkina T, Mani DR, Xu C, Guajardo R, Xie Q, Li T, et al. 2020. Cell-Surface Proteomic Profiling in the Fly Brain Uncovers Wiring Regulators. Cell. 180(2):373–386.e15.

Lo Giudice Q, Leleu M, La Manno G, Fabre PJ. 2019. Single-cell transcriptional logic of cell-fate specification and axon guidance in early-born retinal neurons. Development. 146(17). doi:10.1242/dev.178103. http://dx.doi.org/10.1242/dev.178103.

Love MI, Huber W, Anders S. 2014. Moderated estimation of fold change and dispersion for RNA-seq data with DESeq2. Genome Biol. 15(12):550.

Lyu C, Li Z, Xu C, Kalai J, Luo L. 2026. Rewiring an olfactory circuit by altering cell-surface combinatorial code. Nature. 649(8097):677–684.

Malone JH, Hawkins DL Jr, Michalak P. 2006. Sex-biased gene expression in a ZW sex determination system. J Mol Evol. 63(4):427–436.

Mank JE. 2017. The transcriptional architecture of phenotypic dimorphism. Nat Ecol Evol. 1(1):6.

Mank JE, Nam K, Brunström B, Ellegren H. 2010. Ontogenetic complexity of sexual dimorphism and sex-specific selection. Mol Biol Evol. 27(7):1570–1578.

Mank JE, Rideout EJ. 2021. Developmental mechanisms of sex differences: from cells to organisms. Development. 148(19). doi:10.1242/dev.199750. http://dx.doi.org/10.1242/dev.199750.

Manoli DS, Foss M, Villella A, Taylor BJ, Hall JC, Baker BS. 2005. Male-specific fruitless specifies the neural substrates of Drosophila courtship behaviour. Nature. 436(7049):395–400.

Markow TA, O’Grady PM. 2005. Evolutionary genetics of reproductive behavior in Drosophila: connecting the dots. Annu Rev Genet. 39:263–291.

Marti-Solano M. 2023. A multi-dimensional view of context-dependent G protein-coupled receptor function. Biochem Soc Trans. 51(1):13–20.

McLaughlin CN, Brbić M, Xie Q, Li T, Horns F, Kolluru SS, Kebschull JM, Vacek D, Xie A, Li J, et al. 2021. Single-cell transcriptomes of developing and adult olfactory receptor neurons in. Elife. 10. doi:10.7554/eLife.63856. http://dx.doi.org/10.7554/eLife.63856.

Meissner GW, Luo SD, Dias BG, Texada MJ, Baker BS. 2016. Sex-specific regulation of Lgr3 in Drosophila neurons. Proc Natl Acad Sci U S A. 113(9):E1256–65.

Michael V, Goffinet J, Pearson J, Wang F, Tschida K, Mooney R. 2020. Circuit and synaptic organization of forebrain-to-midbrain pathways that promote and suppress vocalization. Elife. 9. doi:10.7554/eLife.63493. http://dx.doi.org/10.7554/eLife.63493.

Müller L, Grath S, von Heckel K, Parsch J. 2012. Inter- and intraspecific variation in Drosophila genes with sex-biased expression. Int J Evol Biol. 2012:963976.

Naidu VG, Zhang Y, Lowe S, Ray A, Zhu H, Li X. 2020. Temporal progression of Drosophila medulla neuroblasts generates the transcription factor combination to control T1 neuron morphogenesis. Dev Biol. 464(1):35–44.

Naqvi S, Godfrey AK, Hughes JF, Goodheart ML, Mitchell RN, Page DC. 2019. Conservation, acquisition, and functional impact of sex-biased gene expression in mammals. Science. 365(6450). doi:10.1126/science.aaw7317. http://dx.doi.org/10.1126/science.aaw7317.

Nojima T, Rings A, Allen AM, Otto N, Verschut TA, Billeter J-C, Neville MC, Goodwin SF. 2021. A sex-specific switch between visual and olfactory inputs underlies adaptive sex differences in behavior. Curr Biol. 31(6):1175–1191.e6.

Özel MN, Simon F, Jafari S, Holguera I, Chen Y-C, Benhra N, El-Danaf RN, Kapuralin K, Malin JA, Konstantinides N, et al. 2021. Neuronal diversity and convergence in a visual system developmental atlas. Nature. 589(7840):88–95.

Palmateer CM, Artikis C, Brovero SG, Friedman B, Gresham A, Arbeitman MN. 2023. Single-cell transcriptome profiles of -expressing neurons from both sexes. Elife. 12. doi:10.7554/eLife.78511. http://dx.doi.org/10.7554/eLife.78511.

Pan Y, Peng Q, Chen D, Wang R, Xing L, Li C, Ma M, Guan W, Han J, Sun Y. 2025. A master gene for Drosophila male courtship prevents the behavior in females via a truncated protein. Research Square. doi:10.21203/rs.3.rs-6869543/v1. https://www.researchsquare.com/article/rs-6869543/v1.

Paradis E, Schliep K. 2019. ape 5.0: an environment for modern phylogenetics and evolutionary analyses in R. Bioinformatics. 35(3):526–528.

Pavlou HJ, Goodwin SF. 2013. Courtship behavior in Drosophila melanogaster: towards a “courtship connectome.” Curr Opin Neurobiol. 23(1):76–83.

von Philipsborn AC, Liu T, Yu JY, Masser C, Bidaye SS, Dickson BJ. 2011. Neuronal control of Drosophila courtship song. Neuron. 69(3):509–522.

Ranz JM, Castillo-Davis CI, Meiklejohn CD, Hartl DL. 2003. Sex-dependent gene expression and evolution of the Drosophila transcriptome. Science. 300(5626):1742–1745.

Revell LJ. 2024. phytools 2.0: an updated R ecosystem for phylogenetic comparative methods (and other things). PeerJ. 12:e16505.

Rezával C, Pattnaik S, Pavlou HJ, Nojima T, Brüggemeier B, D’Souza LAD, Dweck HKM, Goodwin SF. 2016. Activation of Latent Courtship Circuitry in the Brain of Drosophila Females Induces Male-like Behaviors. Curr Biol. 26(18):2508–2515.

Rice WR. 1984. SEX CHROMOSOMES AND THE EVOLUTION OF SEXUAL DIMORPHISM. Evolution. 38(4):735–742.

Rice WR, Chippindale AK. 2001. Intersexual ontogenetic conflict. J Evol Biol. 14(5):685–693.

Rodríguez-Montes L, Ovchinnikova S, Yuan X, Studer T, Sarropoulos I, Anders S, Kaessmann H, Cardoso-Moreira M. 2023. Sex-biased gene expression across mammalian organ development and evolution. Science. 382(6670):eadf1046.

Ryner LC, Goodwin SF, Castrillon DH, Anand A, Villella A, Baker BS, Hall JC, Taylor BJ, Wasserman SA. 1996. Control of male sexual behavior and sexual orientation in Drosophila by the fruitless gene. Cell. 87(6):1079–1089.

Siwicki KK, Kravitz EA. 2009. Fruitless, doublesex and the genetics of social behavior in Drosophila melanogaster. Curr Opin Neurobiol. 19(2):200–206.

Small CM, Carney GE, Mo Q, Vannucci M, Jones AG. 2009. A microarray analysis of sex- and gonad-biased gene expression in the zebrafish: evidence for masculinization of the transcriptome. BMC Genomics. 10:579.

Stewart AD, Pischedda A, Rice WR. 2010. Resolving intralocus sexual conflict: genetic mechanisms and time frame. J Hered. 101 Suppl 1(Suppl 1):S94–9.

Stockinger P, Kvitsiani D, Rotkopf S, Tirián L, Dickson BJ. 2005. Neural circuitry that governs Drosophila male courtship behavior. Cell. 121(5):795–807.

Stürner T, Brooks P, Serratosa Capdevila L, Morris BJ, Javier A, Fang S, Gkantia M, Cachero S, Beckett IR, Marin EC, et al. 2025. Comparative connectomics of Drosophila descending and ascending neurons. Nature. 643(8070):158–172.

Suvorov A, Kim BY, Wang J, Armstrong EE, Peede D, D’Agostino ERR, Price DK, Waddell P, Lang M, Courtier-Orgogozo V, et al. 2022. Widespread introgression across a phylogeny of 155 Drosophila genomes. Curr Biol. 32(1):111–123.e5.

Sylvestre F, Aubin-Horth N, Bernatchez L. 2025. Sex-biased gene expression across tissues reveals unexpected differentiation in the gills of the threespine stickleback. Peer Community J. 5(e6). doi:10.24072/pcjournal.507. https://peercommunityjournal.org/articles/10.24072/pcjournal.507/.

Tamura K, Subramanian S, Kumar S. 2004. Temporal patterns of fruit fly (Drosophila) evolution revealed by mutation clocks. Mol Biol Evol. 21(1):36–44.

Toker IA, Ripoll-Sánchez L, Geiger LT, Sussfeld A, Saini KS, Beets I, Vértes PE, Schafer WR, Ben-David E, Hobert O. 2025. Divergence in neuronal signaling pathways despite conserved neuronal identity among Caenorhabditis species. Curr Biol. 35(12):2927–2945.e7.

Tosto NM, Beasley ER, Wong BBM, Mank JE, Flanagan SP. 2023. The roles of sexual selection and sexual conflict in shaping patterns of genome and transcriptome variation. Nat Ecol Evol. 7(7):981–993.

Walsh JT, Junker IP, Chen Y-CD, Chen Y-C, Gifford H, Chen DS, Ding Y. 2025. High-resolution single-cell analyses reveal evolutionary constraints and evolvability of sexual circuits in. Proc Natl Acad Sci U S A. 122(47):e2516083122.

Wang F, Wang K, Forknall N, Patrick C, Yang T, Parekh R, Bock D, Dickson BJ. 2020. Neural circuitry linking mating and egg laying in Drosophila females. Nature. 579(7797):101–105.

Wang K, Wang F, Forknall N, Yang T, Patrick C, Parekh R, Dickson BJ. 2021. Neural circuit mechanisms of sexual receptivity in Drosophila females. Nature. 589(7843):577–581.

Wang X, Terfve C, Rose JC, Markowetz F. 2011. HTSanalyzeR: an R/Bioconductor package for integrated network analysis of high-throughput screens. Bioinformatics. 27(6):879–880.

Wayne ML, Telonis-Scott M, Bono LM, Harshman L, Kopp A, Nuzhdin SV, McIntyre LM. 2007. Simpler mode of inheritance of transcriptional variation in male Drosophila melanogaster. Proc Natl Acad Sci U S A. 104(47):18577–18582.

Wei Y-C, Wang S-R, Jiao Z-L, Zhang W, Lin J-K, Li X-Y, Li S-S, Zhang X, Xu X-H. 2018. Medial preoptic area in mice is capable of mediating sexually dimorphic behaviors regardless of gender. Nat Commun. 9(1):279.

Werling DM, Parikshak NN, Geschwind DH. 2016. Gene expression in human brain implicates sexually dimorphic pathways in autism spectrum disorders. Nature Communications. 7(1):1–11.

Wickham H, Averick M, Bryan J, Chang W, McGowan L, François R, Grolemund G, Hayes A, Henry L, Hester J, et al. 2019. Welcome to the tidyverse. J Open Source Softw. 4(43):1686.

Williams TM, Selegue JE, Werner T, Gompel N, Kopp A, Carroll SB. 2008. The regulation and evolution of a genetic switch controlling sexually dimorphic traits in Drosophila. Cell. 134(4):610–623.

Wu T, Hu E, Xu S, Chen M, Guo P, Dai Z, Feng T, Zhou L, Tang W, Zhan L, et al. 2021. clusterProfiler 4.0: A universal enrichment tool for interpreting omics data. Innovation (Camb). 2(3):100141.

Xie C, Künzel S, Tautz D. 2024. Fast evolutionary turnover and overlapping variances of sex-biased gene expression patterns defy a simple binary classification of sexes. eLife. 13. doi:10.7554/eLife.99602.1. [accessed 2025 May 8]. https://elifesciences.org/reviewed-preprints/99602.

Xie C, Künzel S, Tautz D. 2025. Fast evolutionary turnover and overlapping variances of sex-biased gene expression patterns defy a simple binary sex classification of somatic tissues. Elife. 13. doi:10.7554/eLife.99602. http://dx.doi.org/10.7554/eLife.99602.

Yamamoto D, Koganezawa M. 2013. Genes and circuits of courtship behaviour in Drosophila males. Nat Rev Neurosci. 14(10):681–692.

Yang X, Schadt EE, Wang S, Wang H, Arnold AP, Ingram-Drake L, Drake TA, Lusis AJ. 2006. Tissue-specific expression and regulation of sexually dimorphic genes in mice. Genome Res. 16(8):995–1004.

Yapici N, Kim Y-J, Ribeiro C, Dickson BJ. 2008. A receptor that mediates the post-mating switch in Drosophila reproductive behaviour. Nature. 451(7174):33–37.

Ye D, Walsh JT, Junker IP, Ding Y. 2024. Changes in the cellular makeup of motor patterning circuits drive courtship song evolution in Drosophila. Curr Biol. 34(11):2319–2329.e6.

Zappia L, Oshlack A. 2018. Clustering trees: a visualization for evaluating clusterings at multiple resolutions. Gigascience. 7(7). doi:10.1093/gigascience/giy083. http://dx.doi.org/10.1093/gigascience/giy083.

Zhang Y, Sturgill D, Parisi M, Kumar S, Oliver B. 2007. Constraint and turnover in sex-biased gene expression in the genus Drosophila. Nature. 450(7167):233–237.

Zhou C, Pan Y, Robinett CC, Meissner GW, Baker BS. 2014. Central brain neurons expressing doublesex regulate female receptivity in Drosophila. Neuron. 83(1):149–163.

Zhu A, Ibrahim JG, Love MI. 2019. Heavy-tailed prior distributions for sequence count data: removing the noise and preserving large differences. Bioinformatics. 35(12):2084–2092.

