## Supplementary Figure 1 for "Organization and evolution of sex-biased gene expression in *Drosophila* adult sexual circuits"

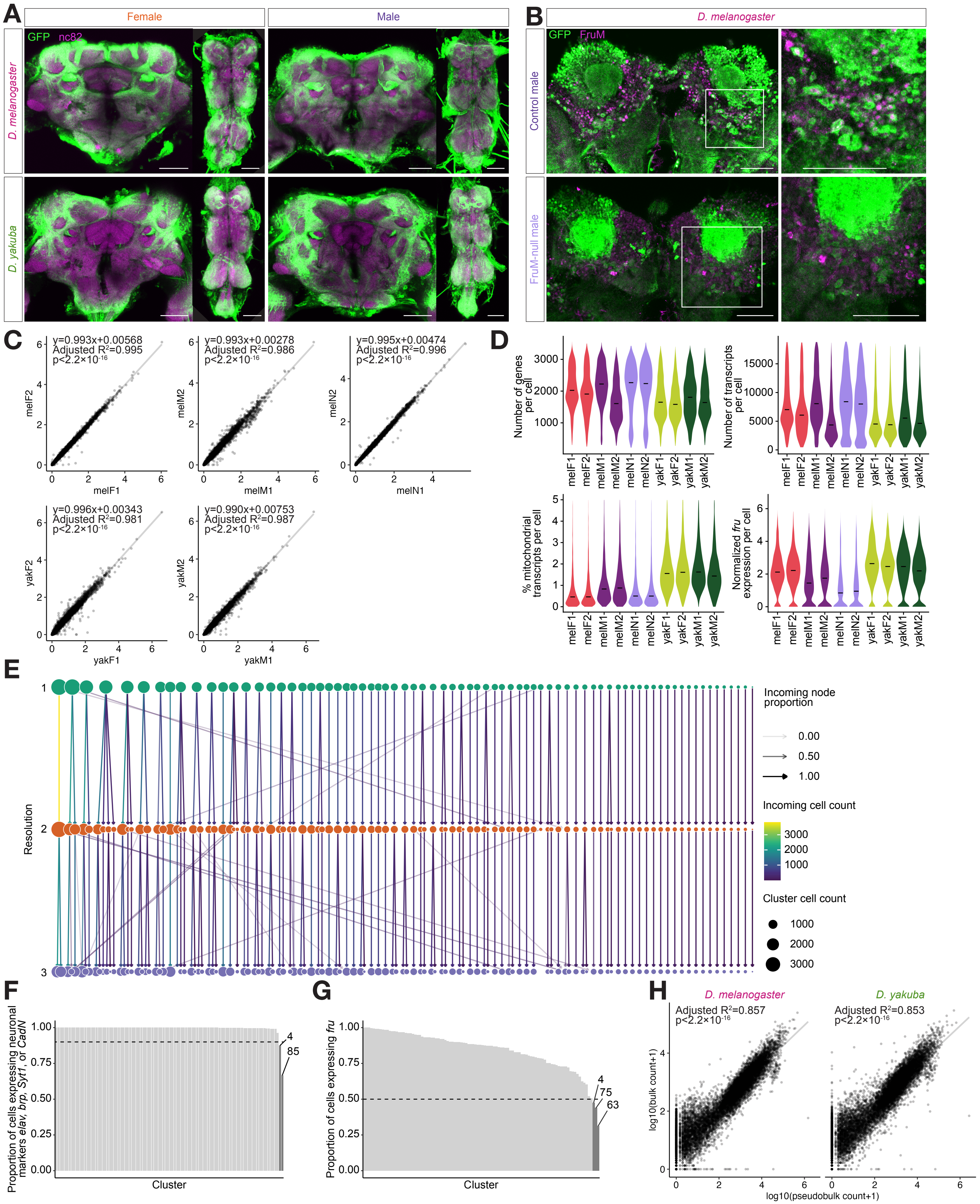

A
Female
Male
GFP nc82
D. melanogaster
D. yakuba
B
D. melanogaster
GFP FruM
Control male
FruM-null male
C
y=0.993x+0.00568
Adjusted R²=0.995
p<2.2×10-16
melF2
melF1
y=0.993x+0.00278
Adjusted R²=0.986
p<2.2×10-16
melM2
melM1
y=0.995x+0.00474
Adjusted R²=0.996
p<2.2×10-16
melN2
melN1
y=0.996x+0.00343
Adjusted R²=0.981
p<2.2×10-16
yakF2
yakF1
y=0.990x+0.00753
Adjusted R²=0.987
p<2.2×10-16
yakM2
yakM1
D
Number of genes per cell
Number of transcripts per cell
% mitochondrial transcripts per cell
Normalized fru expression per cell
melF1 melF2 melM1 melM2 melN1 melN2 yakF1 yakF2 yakM1 yakM2
E
Resolution
Incoming node proportion
0.00
0.50
1.00
Incoming cell count
3000
2000
1000
Cluster cell count
1000
2000
3000
F
Proportion of cells expressing neuronal markers elav, brp, Syt1, or CadN
Cluster
4
85
G
Proportion of cells expressing fru
Cluster
4
75
63
H
D. melanogaster
Adjusted R²=0.857
p<2.2×10-16
D. yakuba
Adjusted R²=0.853
p<2.2×10-16
log10(bulk count+1)
log10(pseudobulk count+1)

**Figure S1 | *fru-GAL4* reagents and sample quality metrics.**
**(A)** Confocal images of *fru-GAL4/UAS-myrGFP* females and males in *D. melanogaster* and *D. yakuba*. Brain images (left) show maximum projection of optic slices that contain *fru* pC1 neurons, and VNC images (right) show maximum projection of the ventral side of VNC. Scale bars: 50 μm.
**(B)** Confocal images of *D. melanogaster UAS-mCD8::GFP/+; fru-GAL4/+* control males and *UAS-mCD8::GFP/+; fru-GAL4/fru-GAL4* FruM null males. The white square in the left panel illustrates the region of the brain shown in the other panels. The anti-FruM antibody exhibits non-specific binding. Scale bars: 50 μm.
**(C)** Correlation of log-normalized gene expression values of all genes between pseudobulked replicates of the same sample type.
**(D)** Violin plots showing the distributions of gene count, transcript count, mitochondrial transcripts percentage, and log-normalized *fru* expression across all cells in each sample type.
**(E)** Relationships between clusters defined at resolution=1 (teal), 2 (orange), and 3 (purple).
**(F)** Proportion of cells in each cluster expressing any of the neuronal marker genes *elav*, *brp, Syt1,* or *CadN*. The dotted horizontal line denotes the cutoff at 0.9. Dark gray bars represent clusters with <90% of cells expressing any of the neuronal markers, and are removed from subsequent analyses.
**(G)** Proportion of cells in each cluster expressing *fru*. The dotted horizontal line denotes the cutoff at 0.5. Dark gray bars represent clusters with <50% *fru*+ cells, and are removed from subsequent analyses.
**(H)** Correlation of log-transformed transcript counts (+1) in pseudobulked single-cell RNA-seq data *versus* bulk RNA-seq data in *D. melanogaster* and *D. yakuba* males.
Dmel: *D. melanogaster*; Dyak: *D. yakuba*; F: female; M: male; N: FruM-null male.
