## Supplementary Figure 2 for "Organization and evolution of sex-biased gene expression in *Drosophila* adult sexual circuits"

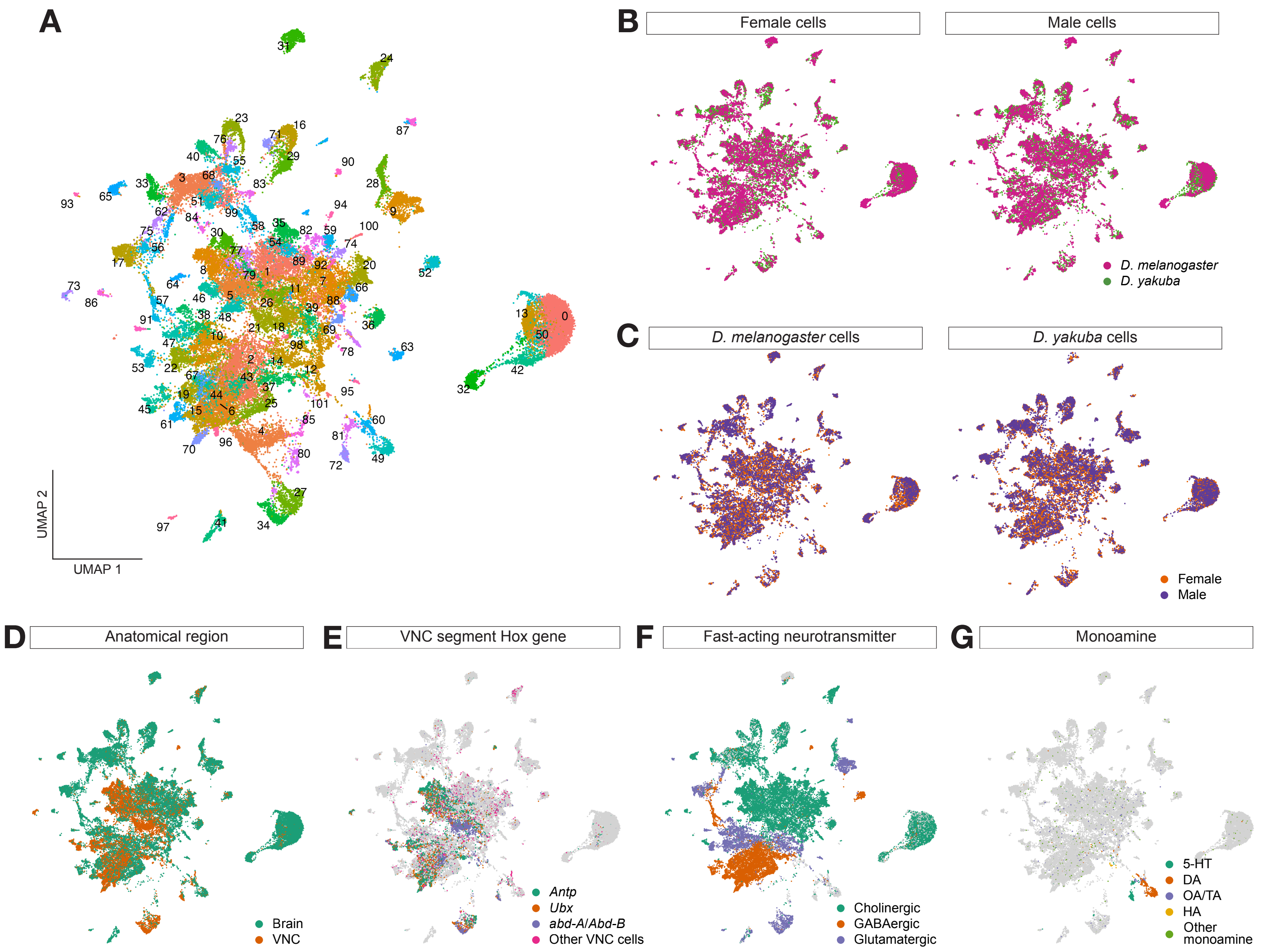


**Figure S2 | Sample integration and *fru* neurons annotation.**

**(A)** Uniform Manifold Approximation and Projection (UMAP) of all cells in the dataset, color-coded by and labeled with cluster identity.

**(B)** UMAP of female and male cells colored by species: *D. melanogaster* cells are shown in magenta and *D. yakuba* cells are shown in green.

**(C)** UMAP of *D. melanogaster* and *D. yakuba* cells colored by sex: female cells are shown in orange and male cells are shown in purple.

**(D)** UMAP of cells colored by anatomical region (head or ventral nerve cord (VNC)) based on marker gene expression: VNC is annotated by the expression of *Antp*, *Ubx*, *adb-A,* or *Abd-B,* or a higher expression of *tsh* than *oc*; head is annotated by the expression of *oc* or one of the neuronal markers *elav, brp, Syt1*, or *CadN* in non-VNC cells.

**(E)** UMAP of cells colored by their most highly expressed VNC segment Hox genes among *Antp*, *Ubx*, or *adb-A*/*Abd-B*.

**(F)** UMAP of cells colored by their fast-acting neurotransmitter identity.

**(G)** UMAP of cells colored by monoamine identity. 5-HT: serotonin; DA: dopamine; OA: octopamine; TA: tyramine; HA: histamine.

In **(B-G)**, the four low confidence clusters and FruM-null samples are not shown.
