## Supplementary Figure 3 for "Organization and evolution of sex-biased gene expression in *Drosophila* adult sexual circuits"

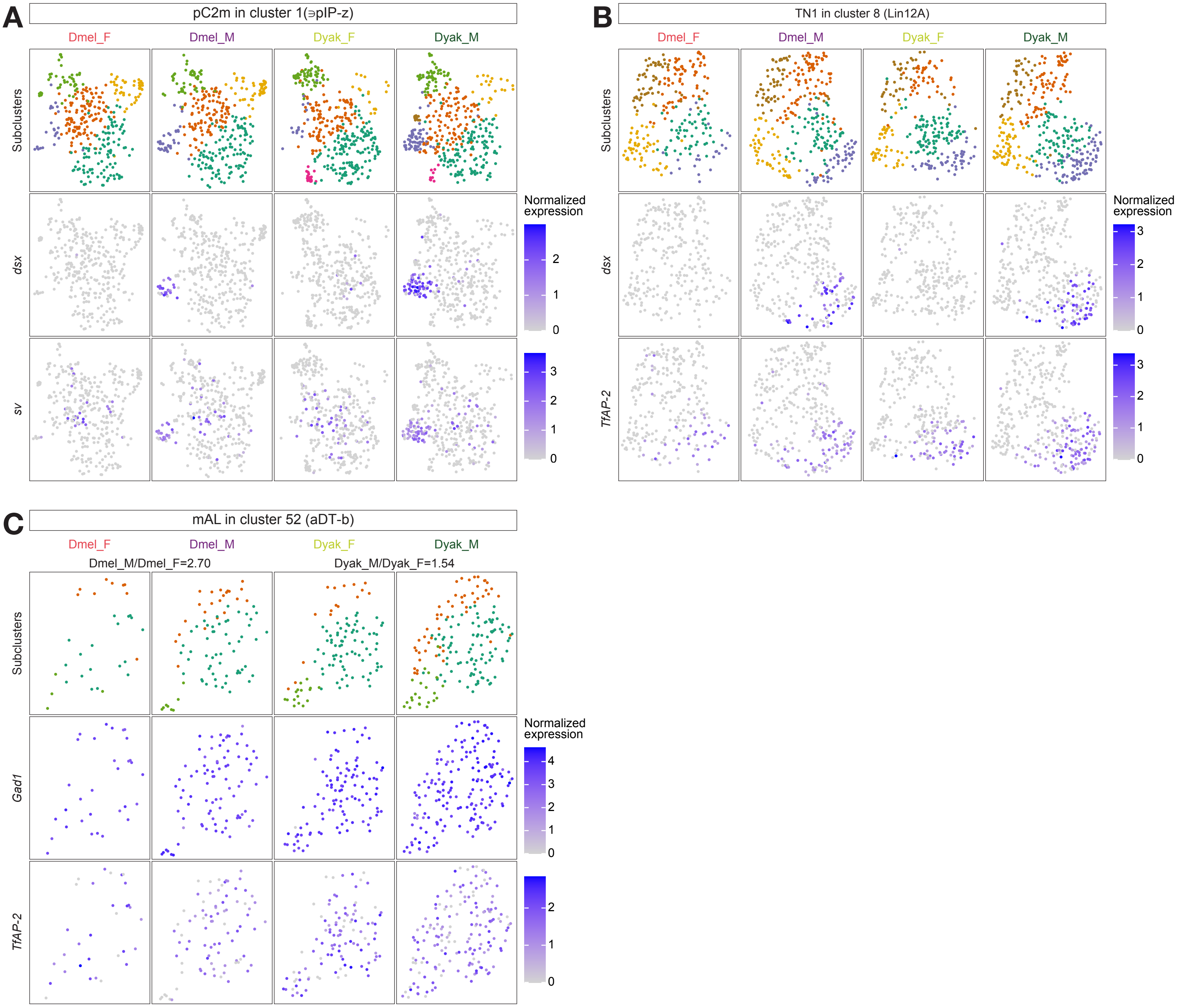


**Figure S3 | Populations with male-biased abundance.**

**(A)** Cluster 1 plotted in its own Uniform Manifold Approximation and Projection (UMAP) space, separated by sample type. The top row shows cells color-coded by their subcluster identity. The middle and bottom rows show cells color-coded by normalized expression of pC2m marker genes *dsx* and *sv*, respectively. Color scales for their respective gene are shown to the right.

**(B)** Cluster 8 plotted in its own UMAP space, separated by sample type. The top row shows cells color-coded by their subcluster identity. The middle and bottom rows show cells color-coded by normalized expression of TN1 marker genes *dsx* and *TfAP-2*, respectively. Color scales for their respective gene are shown to the right.

**(C)** Cluster 52 plotted in its own UMAP space, separated by sample type. The ratios of normalized cluster 52 cells in males to that in females is shown above the top row. The top row shows cells color-coded by their subcluster identity. The middle and bottom rows show cells color-coded by normalized expression of mAL marker genes *Gad1* and *TfAP-2*, respectively. Color scales for their respective gene are shown to the right. Dmel_F: *D. melanogaster* female; Dmel_M: *D. melanogaster* male; Dyak_F: *D. yakuba* female; Dyak_M: *D. yakuba* male.
