## Supplementary Figure 4 for "Organization and evolution of sex-biased gene expression in *Drosophila* adult sexual circuits"

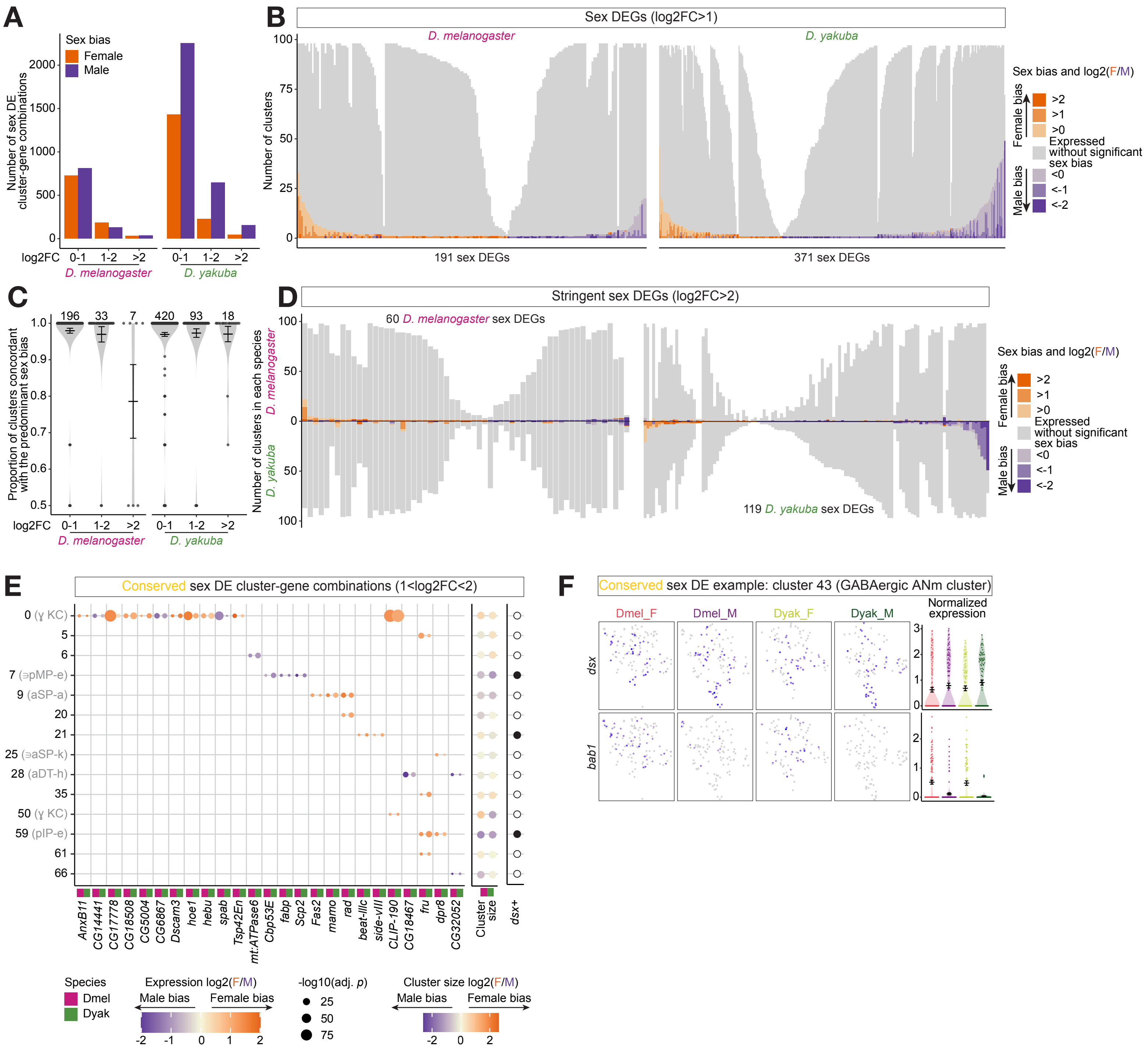


**Figure S4 | Sex differences are limited and largely species-specific.**

**(A)** Number of sex differentially expressed (DE) cluster-gene combinations in at different thresholds of log2-transformed ratio of aggregated female to male gene expression (log2FC) in each species, color-coded by sex bias.

**(B)** Number of clusters in which each *D. melanogaster* (left) and *D. yakuba* (right) sex DE genes (DEG) is expressed and sex DE in its respective species. Clusters in which the sex DEG is expressed without significant sex-bias are shown in gray, and clusters with sex-biased expression are shown in shades of orange (female-biased) or purple (male-biased) based on their log2FC.

**(C)** Concordance of the direction of sex bias, measured as the proportion of clusters that are concordant with the predominant sex bias, of sex DEGs that are DE in more than one cluster defined at different log2FC thresholds. Concordance values range between 0.5 (half of the DE clusters are biased in each direction) and 1 (all DE clusters have the same sex bias). Number of genes in each group are shown at the top.

**(D)** Comparison of *D. melanogaster* (left) and *D. yakuba* (right) sex DEG expression in *D. melanogaster* (above x-axis) *versus D. yakuba* (below x-axis). Clusters in which the sexDEG is expressed without significant sex bias are shown in gray, and clusters with sex-biased expression are shown in shades of orange (female-biased) or purple (male-biased) based on their log2FC.

**(E)** Log2FC and -log10-transformed adjusted p-value (-log10(adj. p)) of conserved sex DE cluster-gene combinations in both species. Log2-transformed ratio of female cells to male cells and *dsx* expression status for each cluster are shown to the right. Size scale of -log10(adj. p) and color scales of log2FC are shown next to the plot. Magenta and green rectangles in the margin of each column denote *D. melanogaster* and *D. yakuba*, respectively.

**(F)** Cluster 43 plotted in its own UMAP space, separated by sample type. The top and bottom rows show cells color-coded by normalized *dsx* and *bab1* expression, respectively. Normalized expression of each gene in each cell, grouped by sample type, are shown to the right. Error bars show mean±SEM.

DEG: differentially expressed gene; ANm: abdominal neuromere; Dmel: *D. melanogaster*; Dyak: *D. yakuba;* F: female; M: male.
