## Supplementary Figure 5 for "Organization and evolution of sex-biased gene expression in *Drosophila* adult sexual circuits"

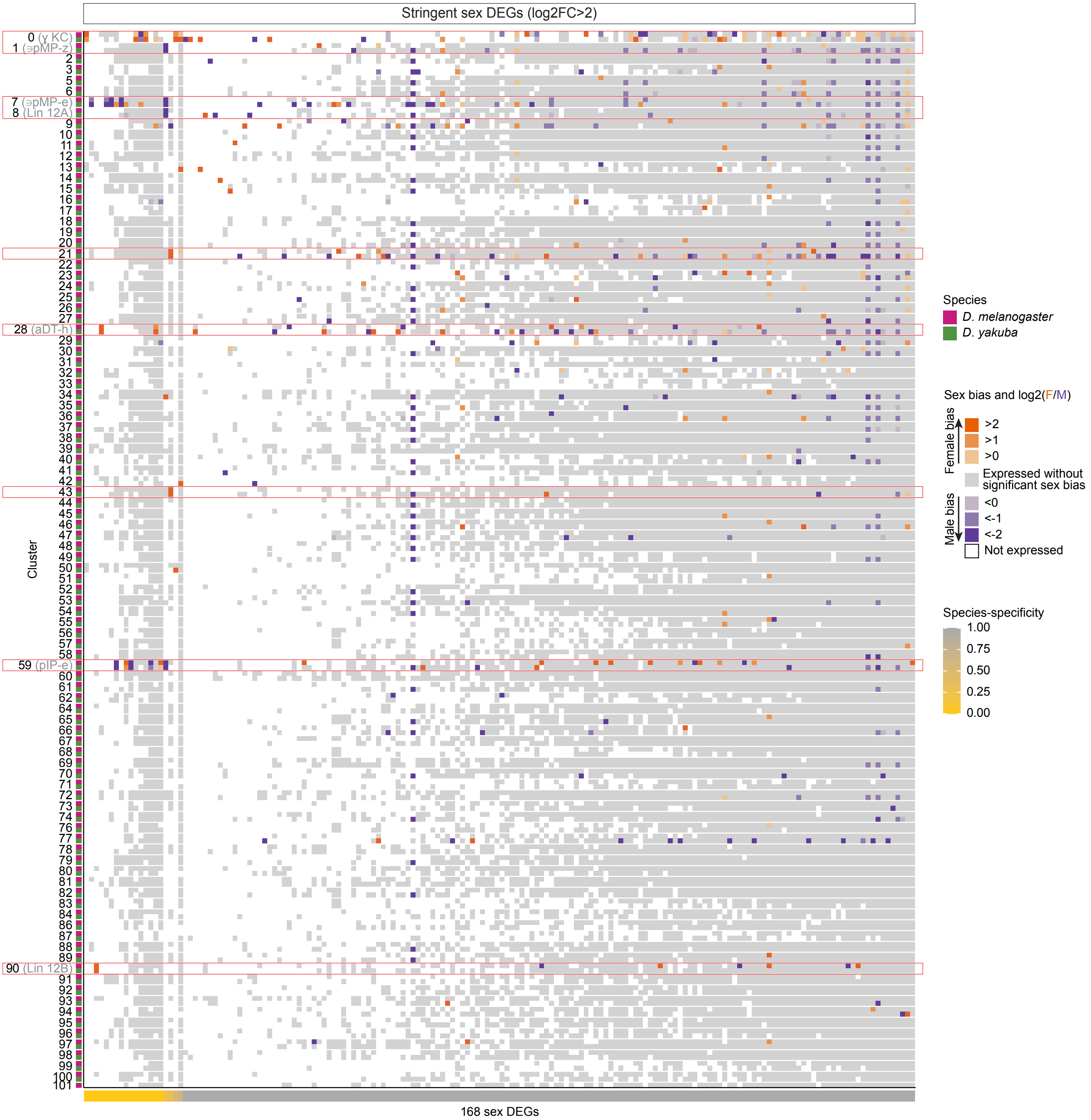


**Figure S5 | Sex differentially expressed genes are localized and largely species-specific.**
Expression and sex bias of all sex differentially expressed genes (DEGs) at the stringent threshold (log2-transformed ratio of aggregated female to male gene expression (log2FC)>2). Clusters with sex-biased expression are shown in shades of orange (female-biased) or purple (male-biased) based on their log2FC, clusters in which the sex DEG are expressed without significant sex-bias are shown in gray, and clusters in which the sex DEG are not expressed are shown in white. Magenta and green rectangles in the margins of each row denote *D. melanogaster* and *D. yakuba* data, respectively. Gradient in the margins of each column denotes each sex DEG's species-specificity. Clusters with conserved sex DE cluster-gene combinations, as shown in Figure 2H, are highlighted with red outlines.
KC: Kenyon cells; F: female; M: male.
