## Supplementary Figure 6 for "Organization and evolution of sex-biased gene expression in *Drosophila* adult sexual circuits"

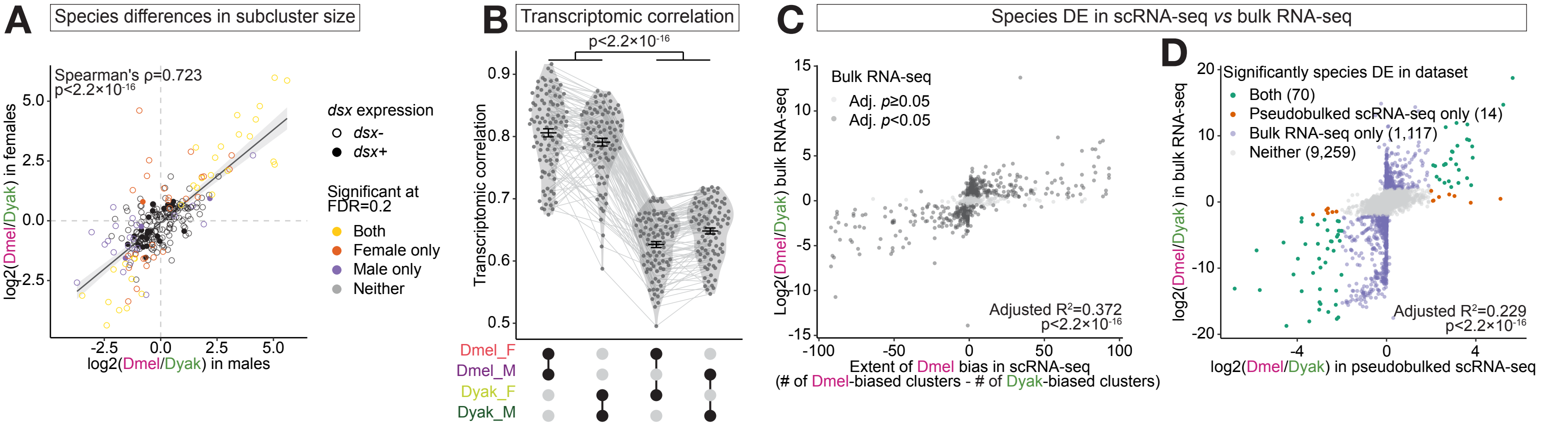


**Figure S6 | Species differences are prevalent and extensively sex-coupled.**

**(A)** Correlation of log2-transformed ratio of *D. melanogaster* cells to *D. yakuba* cells in each subcluster in females versus males. Filled dots denote *dsx+* clusters, and empty dots denote *dsx-* clusters. Dots are color-coded by the sex(es) in which their species bias is statistically significant at FDR=0.2. Regression line and 95% confidence interval are shown behind data points in gray.

**(B)** Inter-sex and inter-species Kendall's tau transcriptomic correlation between pairs of sample types. Error bars show mean±SEM. Light gray lines behind data points link the same cluster in all comparison groups. Statistical significance is tested with ANOVA on a linear model that also accounts for cluster identity, normalized cluster size (log-transformed), inter-replicate Kendall's tau transcriptomic correlation, and number of genes used to calculate transcriptomic correlation.

**(C)** Correlation of each single-cell RNA-seq-defined male species differentially expressed gene (DEG)'s extent of *D. melanogaster* bias against their log2-transformed ratio of *D. melanogaster* to *D. yakuba* gene expression (log2FC) in the bulk RNA-seq data. Extent of *D. melanogaster* bias is calculated as the difference between the number of *D. melanogaster*-biased and *D. yakuba*-biased clusters for each species DEG.

**(D)** Correlation of log2FC of 10,465 genes tested for species DE in the pseudobulked scRNA-seq data and bulk RNA-seq data. Genes are color-coded by the dataset(s) in which they are significantly species DE (log2FC>2 and adjusted p-value<0.05).

Dmel: *D. melanogaster*; Dyak: *D. yakuba;* F: female; M: male.
