## Supplementary Figure 7 for "Organization and evolution of sex-biased gene expression in *Drosophila* adult sexual circuits"

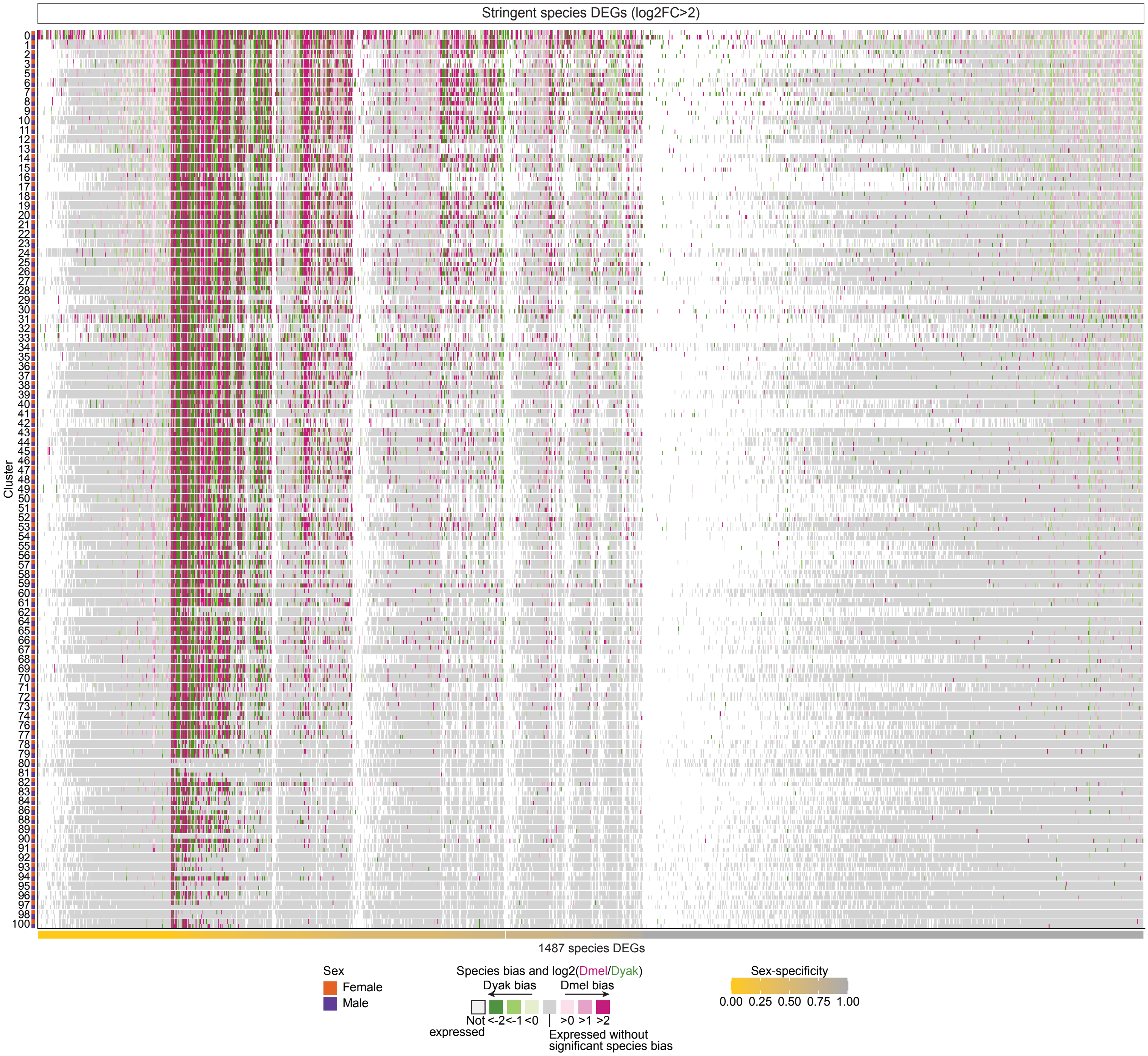


**Figure S7 | Transcriptomic evolution is strongly coupled between sexes.**
Expression and species bias of all species differentially expressed genes (DEGs) at the stringent threshold (log2-transformed ratio of aggregated *D. melanogaster* to *D. yakuba* gene expression (log2FC)>2). Clusters with species-biased expression are shown in shades of magenta (*D. melanogaster*-biased) or green (*D. yakuba*-biased) based on their log2FC, clusters in which the species DEG are expressed without significant species-bias are shown in gray, and clusters in which the species DEG are not expressed are shown in white. Orange and purple rectangles in the margins of each row denote female and male data, respectively. Gradient in the margins of each column denotes each species DEG's sex-specificity.
Dmel: *D. melanogaster*; Dyak: *D. yakuba*.
