## Supplementary Figure 8 for "Organization and evolution of sex-biased gene expression in *Drosophila* adult sexual circuits"

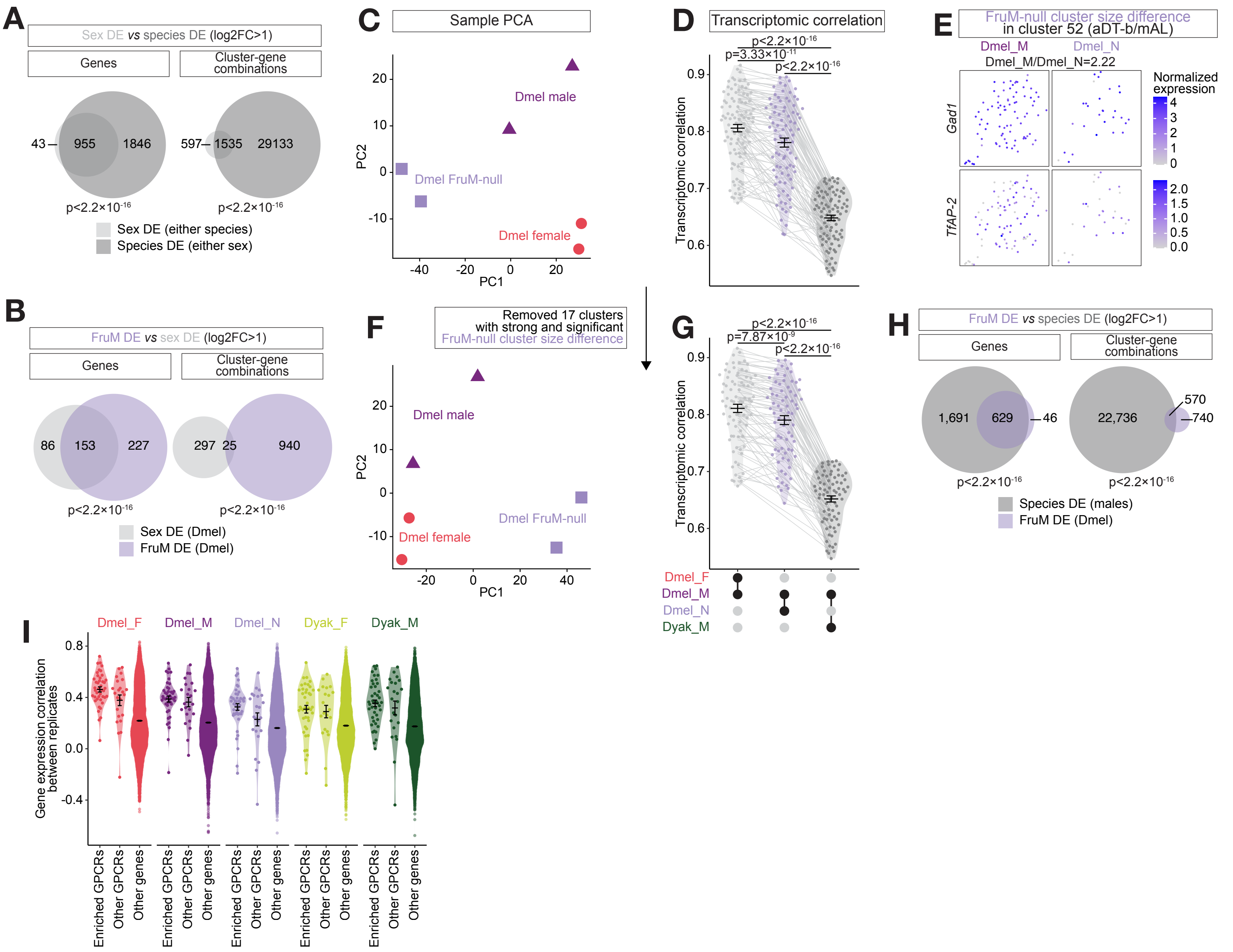


**Figure S8 | Factors shaping gene expression evolution.**

**(A,B,H)** Area-proportional Euler diagrams showing the number of genes and cluster-gene combinations differentially expressed (DE) between sexes (sex DE) and between species (species DE) **(A)**, those DE between sexes in *D. melanogaster* (sex DE) and between FruM-null males and wildtype males (FruM DE) **(B)**, and those DE between species in males (species DE) and between FruM-null males and wildtype males (FruM DE) **(H)**. Statistical significance of the overlap is calculated based on the hypergeometric distribution and shown below each panel.

**(C, F)** Six *D. melanogaster* samples generated in the study along principle components (PCs) 1 and 2, before **(C)** and after **(F)** filtering out the 17 clusters with strong (among the top 20 clusters with the largest fold-change in either direction) and significant (scCODA FDR<0.2) cluster size difference between FruM-null males and control males. PC analysis is performed based on the log-normalized aggregated gene expression of the 2,000 most variable genes across these samples.

**(D, G)** Kendall's tau transcriptomic correlation between sexes in *D. melanogaster*, between wildtype and FruM-null males in *D. melanogaster*, and between species in males, before **(D)** and after **(G)** filtering out the 17 clusters as in **(F)**. Error bars show mean±SEM. Light gray lines behind data points link the same cluster in all comparison groups. Statistical significance is tested with ANOVA on a linear model that also accounts for cluster identity, inter-replicate Kendall's tau transcriptomic correlation, and number of genes used to calculate transcriptomic correlation.

**(E)** Cluster 52 plotted in its own UMAP space, separated by sample type. The top and bottom rows show cells color-coded by normalized *Gad1* and *TfAP-2* expression, respectively.

**(I)** Inter-replicate Kendall's tau transcriptomic correlation of each gene, separated by sample types and by gene groups: G protein-coupled receptors (GPCRs) enriched in the "G protein-coupled receptor activity" term in Figure 4F, other GPCRs, and non-GPCR genes. Error bars show mean±SEM.

Dmel: *D. melanogaster*; Dyak: *D. yakuba*; F: female; M: male; N: FruM-null male.
