## Supplementary Figure 9 for "Organization and evolution of sex-biased gene expression in *Drosophila* adult sexual circuits"

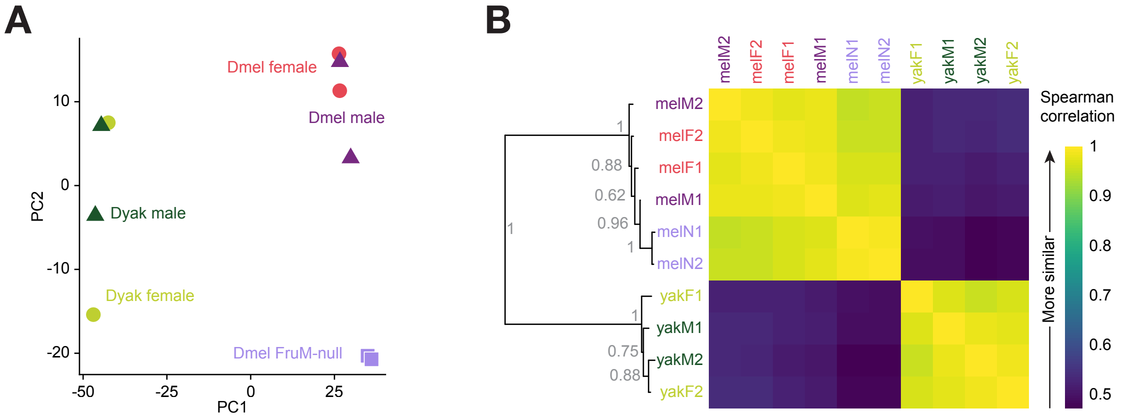


**Figure S9 | Relationships between samples.**
**(A)** Ten samples generated in the study along principle components (PCs) 1 and 2. PC analysis is performed based on the log-normalized aggregated gene expression of the 2,000 most variable genes across the ten samples. Dmel: *D. melanogaster*; Dyak: *D. yakuba.*
**(B)** Left: neighbor-joining tree constructed based on pairwise sample-level Spearman correlation calculated using the log-normalized aggregated gene expression of the 2,000 most variable genes across the ten samples. Numbers next to each node show the proportion of draws (out of 1000 random draws of 1000 genes each, without replacement) that support each branch. Right: Heatmap showing Spearman correlation between each pair of samples. Color scale is shown on the right. mel: *D. melanogaster*; yak: *D. yakuba*; F: female; M: male; N: FruM-null male.
