## Supplementary Figure 10 for "Organization and evolution of sex-biased gene expression in *Drosophila* adult sexual circuits"

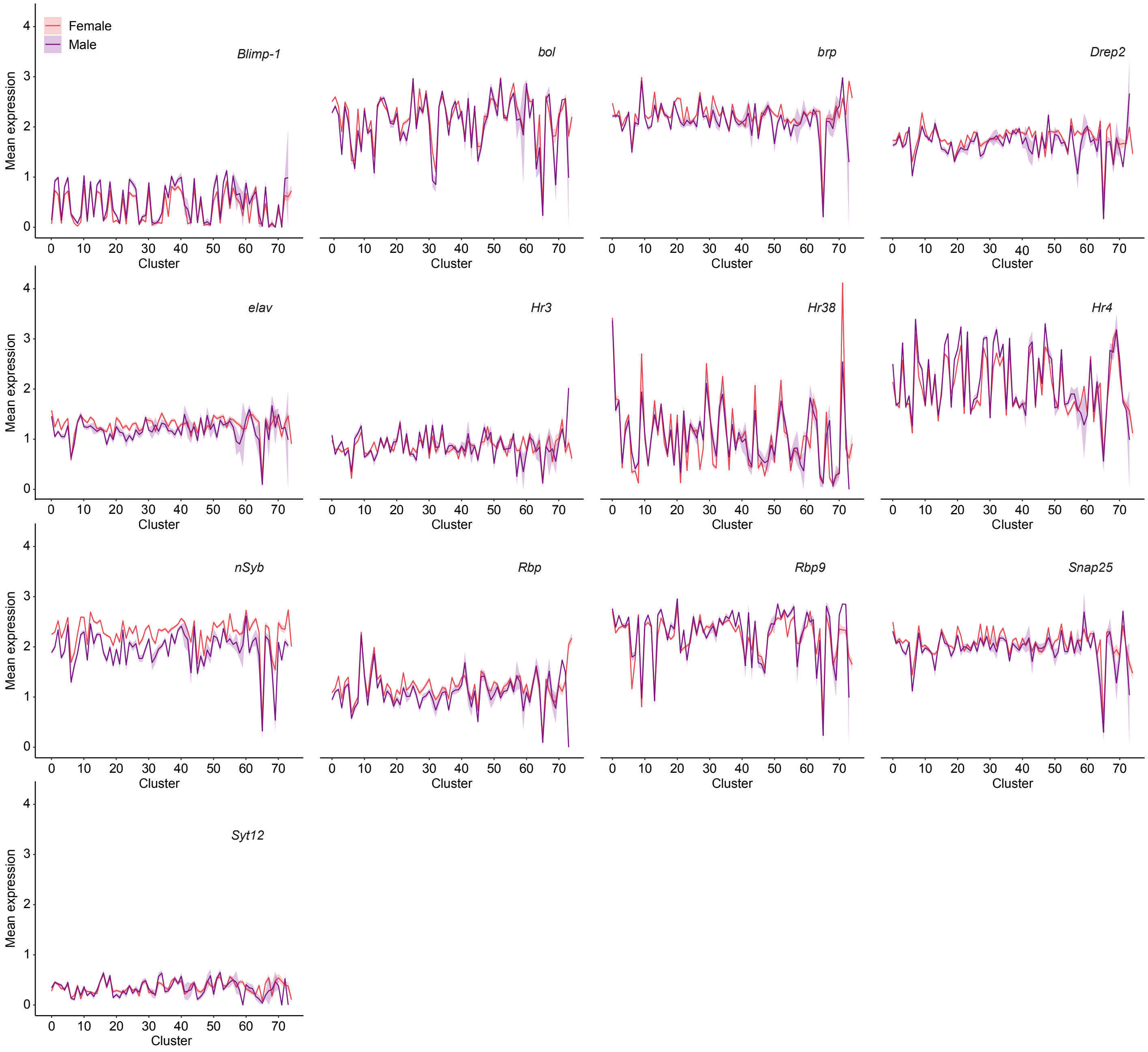


**Figure S10 | Maturation gene expression in pupal data.**
Mean±SEM of normalized gene expression across clusters, showing maturation transcription factors *Hr3* and *Hr4* (Jain et al. 2022, Elkahlah et al. 2025) and 11 additional genes with dynamic and pan-neuronally coordinated expression during pupal development (Kurmangaliyev et al. 2020). Lines are color-coded by sex.
