## Supplementary Table 1 for "Organization and evolution of sex-biased gene expression in *Drosophila* adult sexual circuits"

**Supplementary Table 1 | Sample statistics**

| Sample ID | Number of cells | Median transcript count | Min transcript count | Max transcript count | Median gene count | Min gene count | Max gene count |
|---|---|---|---|---|---|---|---|
| melF1 | 4,901 | 7,017 | 453 | 18,782 | 2,024 | 302 | 3,557 |
| melF2 | 5,430 | 6,062 | 460 | 18,717 | 1,907 | 253 | 3,498 |
| melM1 | 5,741 | 8,056 | 452 | 18,793 | 2,223 | 304 | 3,529 |
| melM2 | 5,224 | 4,348 | 451 | 18,672 | 1,607 | 280 | 3,390 |
| melN1 | 5,133 | 8,403 | 444 | 18,860 | 2,266 | 221 | 3,425 |
| melN2 | 5,903 | 8,000 | 235 | 18,694 | 2,237 | 190 | 3,528 |
| yakF1 | 6,744 | 4,516 | 442 | 18,535 | 1,650 | 252 | 3,421 |
| yakF2 | 5,370 | 4,388 | 461 | 18,088 | 1,583 | 274 | 3,399 |
| yakM1 | 8,264 | 5,543 | 444 | 18,356 | 1,806 | 302 | 3,434 |
| yakM2 | 5,318 | 4,632 | 463 | 18,006 | 1,640 | 301 | 3,359 |
| **Total** | **58,028** | | | | | | |
| **Mean** | **5,802.80** | **6,096.35** | **430.50** | **18,550.30** | **1,894.20** | **267.90** | **3,454.00** |
| **Sd** | **1,006.39** | **1,651.93** | **69.09** | **301.80** | **277.45** | **38.95** | **68.41** |
