## Supplementary Table 2 for "Organization and evolution of sex-biased gene expression in *Drosophila* adult sexual circuits"

**Supplementary Table 2 | Number of neurons, *fru* cells, and *dsx* cells**

| Sample | Number of cells | Number of neurons | Number of *fru* cells | Number of *dsx* cells | Proportion of neurons | Proportion of *fru* cells | Proportion of *dsx* cells |
|---|---|---|---|---|---|---|---|
| Dmel_F | 10,331 | 10,293 | 9,382 | 186 | 0.996 | 0.908 | 0.018 |
| Dmel_M | 10,965 | 10,925 | 8,281 | 422 | 0.996 | 0.755 | 0.038 |
| Dmel_N | 11,036 | 10,920 | 6,902 | 312 | 0.989 | 0.625 | 0.028 |
| Dyak_F | 12,114 | 12,013 | 10,985 | 300 | 0.992 | 0.907 | 0.025 |
| Dyak_M | 13,582 | 13,509 | 11,711 | 754 | 0.995 | 0.862 | 0.056 |
| **Total** | **58,028** | **57,660** | **47,261** | **1,974** | **0.994** | **0.814** | **0.034** |
| **Mean** | **11,605.60** | **11,532.00** | **9,452.20** | **394.80** | **0.994** | **0.812** | **0.033** |
| **Sd** | **1,277.11** | **1,266.69** | **1,957.79** | **217.49** | **0.003** | **0.121** | **0.015** |

Neurons are defined as cells expressing any of the neuronal markers *elav*, *brp*, *Syt1*, or *CadN*.
